# Differential effects of early or late exposure to prenatal maternal immune activation on mouse embryonic neurodevelopment

**DOI:** 10.1101/2021.07.14.452084

**Authors:** Elisa Guma, Maude Bordeleau, Emily Snook, Gabriel Desrosiers-Grégoire, Fernando González Ibáñez, Katherine Picard, Shoshana Spring, Jason P. Lerch, Brian J. Nieman, Gabriel A. Devenyi, Marie-Eve Tremblay, M. Mallar Chakravarty

## Abstract

Exposure to maternal immune activation (MIA) *in utero* is a risk factor for neurodevelopmental and psychiatric disorders. MIA-induced deficits in adolescent and adult offspring have been well characterized, however, less is known about the effects of MIA-exposure on embryo development. To address this gap, we performed high-resolution *ex vivo* magnetic resonance imaging (MRI) to investigate the effects of early (gestational day [GD]9) and late (GD17) MIA-exposure on embryo (GD18) brain structure. We identify striking neuroanatomical changes in the embryo brain, particularly in the late exposed offspring. We further examined hippocampal neuroanatomy using electron microscopy and identified differential effects due to MIA-timing. An increase in apoptotic cell density was observed in the GD9 exposed offspring, while an increase in the density of dark neurons and glia, putative markers for increased neuroinflammation and oxidative stress, was observed in GD17 exposed offspring, particularly in females. Overall, our findings integrate imaging techniques across different scales to identify differential impact of MIA-timing on the earliest stages of neurodevelopment.

## 1. Introduction

Brain development is a remarkable and complex set of processes under the organizational control of genetic, environmental, and immune regulation. The tightly regulated nature and interdependence of these processes make them vulnerable to a variety of risk factors. Converging lines of evidence suggest an association between prenatal exposure to maternal infection and increased risk for a host of neurodevelopmental disorders in offspring, including schizophrenia and autism spectrum disorder (ASD) (1–3). Indeed, exposure to maternal immune activation (MIA) in animal models has been shown to induce neuroanatomical and behavioural changes relevant to many neurodevelopmental disorders (4). MIA leads to an increase in maternal pro-inflammatory cytokines and chemokines, which are thought to interfere with fetal brain development by disturbing its delicate ecosystem, potentially as a consequence of the microglial response (5–9). Exposure to MIA during the sensitive window of *in utero* brain development may alter neurodevelopmental trajectories, thereby increasing risk for neuropsychiatric disorders later in life (10,11). Identifying these sensitive windows and their impact on later development is critical to our understanding of the effects of MIA-exposure. Previous work from our group has demonstrated that the gestational timing of MIA-exposure has a differential impact on offspring brain and behavioural development, with a greater variation observed following early MIA-exposure (gestational day [GD]9) relative to exposure later in gestation (GD17) or to a control (11). These differences may be attributable to variation in maternal immune responsiveness and fetal brain development across gestation (12).

Although there is significant evidence that MIA-exposure *in utero* alters brain development trajectories in both human (13) and animal models (14,15), it is unclear how soon after MIA-exposure these changes can be detected. MIA-induced outcomes have been better characterized in adolescent and adult rodent offspring (12). However, to better understand the initiation and progression of MIA-induced pathology, it is critical to study the impact of MIA-exposure on brain development at the earliest stages of life. Human neuroimaging studies have identified alterations in functional and structural connectivity in the infant brain following exposure to chronic, low-grade inflammation (as measured by interleukin [IL]-6 levels and/or C-reactive protein [CRP] in the maternal plasma) (16–18). Even though some observations in early phases of life have been made, there is less information available regarding how MIA-exposure impacts the morphogenesis of the fetus. Recent work on this topic suggests that MIA-exposure induces acute upregulation of genes involved in immune signaling, hypoxia, and angiogenesis in the fetal mouse brain (19). Further, alterations in neuronal proliferation, neuronal and glial specification, cortical lamination (19,20), global mRNA translation, and altered nascent proteome synthesis have been reported (21).

These findings suggest that effects of MIA-exposure may be detectable in the fetal and neonatal period across mouse and human studies. However, it is unclear whether the transcriptional and histological variation observed in rodents translates to the neuroanatomical changes detected by whole-brain imaging observed in human studies. Furthermore, although gestational timing has been shown to have differential effects in adolescent and adult offspring (11,22), it is unclear how it affects neurodevelopment in its early phase. A better understanding of the neurodevelopmental sequelae of MIA-exposure on very early brain development is of importance in the context of the current COVID-19 pandemic, as mothers who contracted the virus during pregnancy were more likely to have obstetric complications leading to poor fetal health outcomes such as low birth weight, intrauterine growth restriction, and preterm birth (23,24).

To build upon our previous investigations in which we characterized brain development from adolescence to adulthood (11), we aimed to develop a chronology of how the timing of MIA-exposure may impact brain development *in utero*, using the same gestational exposures as in our previous work. We leveraged structural magnetic resonance imaging (MRI), an inherently 3-dimensional imaging technique applicable for mouse phenotyping (25). This technique allows for a comparable assay across species, providing a potential avenue for establishing cross-species homology (26). We examine the effects of *in utero* exposure to early (GD9) or late (GD17) MIA with a viral mimetic, polyinosinic:polycytidylic acid (poly I:C), on embryo brain morphology at GD18 using high-resolution *ex vivo* whole-brain MRI. To better understand the cellular underpinnings of the volumetric changes identified by MRI, we leveraged high-resolution electron microscopy (EM) to examine the density of certain cells including apoptotic cells, dark neurons, and dark glial cells. These dark cells are unique from other neurons and glia as they have distinct ultrastructural characteristics reflective of oxidative stress and are only identifiable with electron microscopy (27). We focused on dark and apoptotic cells as they have been identified as a putative marker for neuroinflammation, cellular stress, apoptosis, and disease in the brain parenchyma (27–29). The dorsal hippocampus was selected as the region of interest as it was differentially affected by GD9 and 17 MIA-timing. Further, alterations in this region have been consistently associated with neuropsychiatric disorders (30,31), as well as in response to MIA-exposure in our previous work (11). Furthermore, an increase in dark cell density has been previously observed in the hippocampus of adult MIA-exposed mouse offspring (32). Our results demonstrate neuroanatomical alterations in the GD18 embryo brain following MIA, with differential effects due to timing in many regions, including the dentate gyrus of the hippocampus. Here, we observed a significant increase in the density of apoptotic cells in the GD9-exposed embryos (but not GD17) relative to the control group, while a qualitative increase in the density of dark cells (both neurons and glia) was observed in the GD17-exposed offspring. These findings suggest that morphological changes due to MIA-exposure are already detectable in the fetal brain, and that the timing of MIA-exposure may differentially impact the brain both at anatomical and cellular levels.

## 2. Materials and methods

### 2.1 Animals, prenatal immune activation, and sample preparation

C57BL/6J female and male mice of breeding age (8-12 weeks old) were subject to timed mating procedures (described in **Supplement 1.1**) to generate pregnant dams. Pregnant dams were randomly assigned to one of four treatment groups (**Figure 1** for experimental design): (1) poly I:C (P1530-25MG polyinosinic:polycytidylic acid sodium salt TLR ligand tested; Sigma Aldrich) (5mg/kg, intraperitoneally) at GD 9 (POL E; 7 embryo dams), (2) 0.9% sterile NaCl solution (saline) at GD 9 (SAL E; 6 embryo dams), (3) poly I:C at GD 17 (5mg/kg, intraperitoneally) (POL L; 7 embryo dams), or (4) saline at GD 17 (SAL L; 4 embryo dams). Additionally, immunostimulatory potential of poly I:C was confirmed in a separate group of dams (**Supplement 1.1** for methods, and **Supplement 2.1** and **Supplementary Table 1** for results).

**Figure 1.**
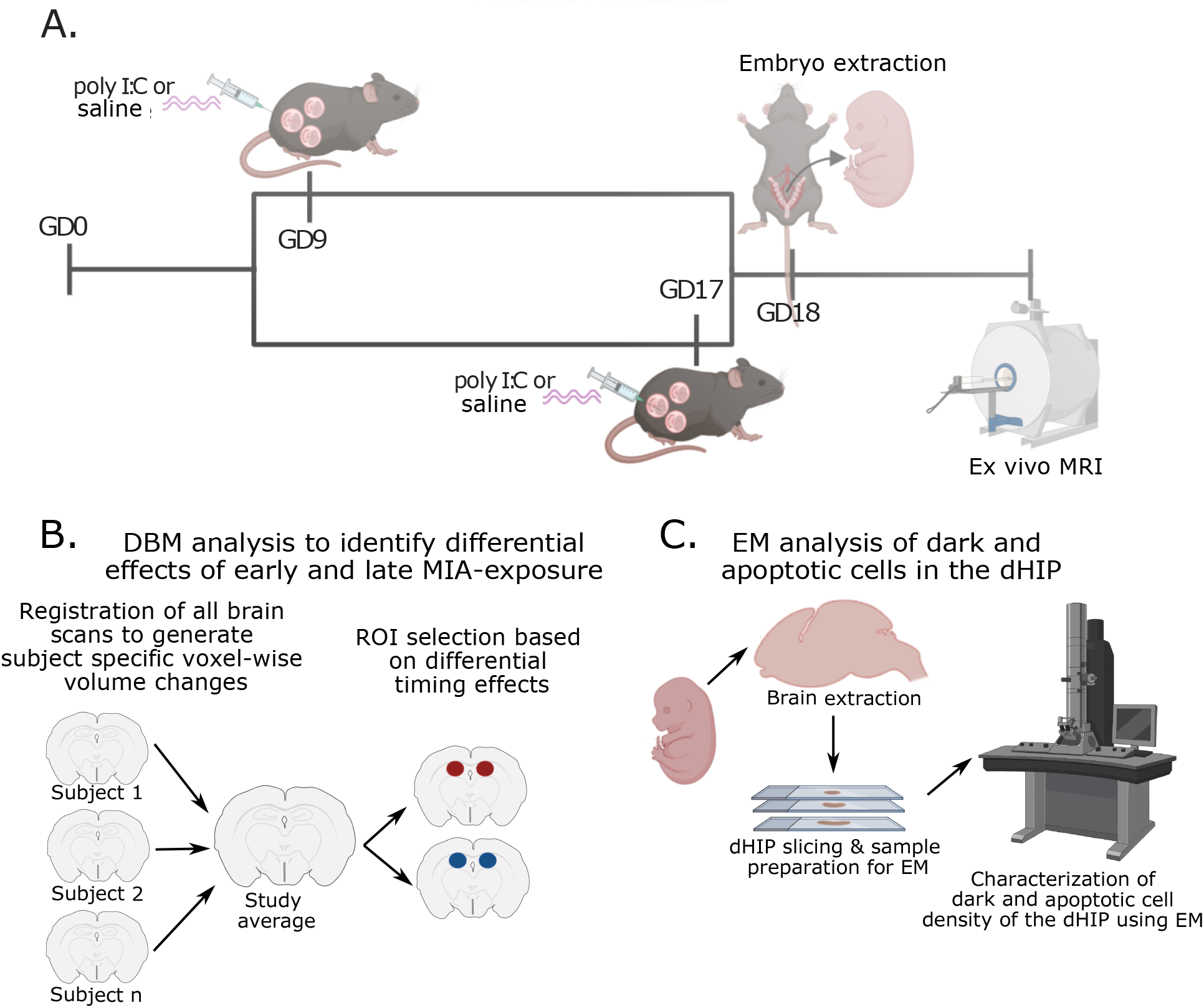
Experimental timeline. **A.** Pregnant dams were injected (i.p.) with poly I:C (5mg/kg) or vehicle (0.9% sterile NaCl) solution on gestational day (GD) 9 or 17. On GD 18, pregnant dams were euthanized, embryos were extracted and prepared for high resolution *ex-vivo* MRI. **B.** Analysis flow of deformation based morphometry analysis used to detect voxel-wise brain volume differences due to early or late MIA-exposure. The dorsal hippocampus was selected as a region of interest for cellular investigation using electron microscopy due to differential effects of timing on bilateral volume. **C.** Embryo brains were extracted from scanned samples, sliced, and prepared for EM investigation of dark glial cell, dark neuron, and apoptotic cell density. DBM: deformation based morphometry; dHIP: dorsal hippocampus; EM: electron microscopy; GD: gestational day; poly I:C: polyinosinic:polycytidylic acid. Figure made using biorender (https://biorender.com).

#### Sample Preparation for MRI

On GD18, pregnant dams were euthanized, embryos were extracted and postfixed in 4% paraformaldehyde (PFA) with 2% gadolinium (MRI contrast agent; Bracco Imaging S.p.A) in PBS for 1 week. A piece of the yolk sac was collected for genotyping of each embryo to identify the sex of the mouse via presence of the SRY gene (performed by Transnetyx, Memphis, TN). Collections were performed in two separate cohorts (with two different poly I:C batches from the same supplier outlined in **Supplement 1.2** and **Table 1**).

**Table 1.** Final sample size for embryo MRI data following quality control.

| Group | Males MRI<br>(Cohort1/Cohort 2) | Females MRI<br>(Cohort1/Cohort 2) | Litters<br>(Cohort1/Cohort 2) |
| --- | --- | --- | --- |
| SAL E | 14 (13/1) | 15 (13/2) | 7 (6/1) |
| SAL L | 12 (10/2) | 12 (6/6) | 4 (2/2) |
| POL E | 11 (6/5) | 17 (9/8) | 7 (4/3) |
| POL L | 14 (11/3) | 17 (15/2) | 7 (6/1) |

### 2.2 Magnetic resonance image acquisition and processing

All samples were shipped to the Mouse Imaging Centre (Toronto, ON) for scanning. A multi-channel 7.0-T MRI scanner with a 40 cm diameter bore (Varian Inc., Palo Alto, CA) was used to acquire anatomical images of the entire embryo (whole body). A custom-built 16-coil solenoid array was used to acquire 40 μm^3^ resolution images from 16 samples concurrently (33) (see **Supplement 1.3.1** for details).

Preprocessed embryo brain images (https://github.com/CoBrALab/documentation/wiki/Embryo-scan-preprocessing) of all subjects in the study were aligned by unbiased deformation based morphometry using the antsMultivariateTemplateConstruction2.sh tool (https://github.com/CoBrALab/twolevel_ants_dbm) (34). The output of this iterative group-wise registration procedure generates a group average from all the scans in the study, as well as the minimum deformation fields that is required to accurately map each individual subject to the average at the voxel level (3D representation of group average in **Supplementary figure 2)**. Relative Jacobian determinants (35), which explicitly model only the non-linear deformations and remove global linear transformation (attributable to differences in total brain size) were blurred using a Gaussian kernel at 160 μm full-width-at-half-maximum to better conform to Gaussian assumptions for downstream statistical testing (see **Supplement 1.3.2** for details).

Collection was performed in two rounds with two different batches of poly I:C (same supplier and product). Number indicates total sample size following quality control. The number of samples coming from each collection cohort is noted in parentheses.

### 2.3 Electron microscopy

After MRI scanning, embryo brains (SAL E, n= 4 [2males/2females]; SAL L, n= 7 [3males/4females]; POL E, n= 8 [4males/4females]; POL L, n= 8 [4males/4females]) were extracted from the fixed samples and further post-fixed with 3.5% acrolein in phosphate buffer [100mM] (pH 7.4) overnight at 4°C. Post-fixed brains were sectioned to 50 μm sagittal slices using a VT1200S vibratome (Leica Biosystem), and stored at −20 °C in cryoprotectant (30% glycerol, 30% ethylene glycol in PBS [50mM] (pH 7.4)). Three brain sections in which the dorsal hippocampus was present (Coronal section 12-15,(36)), roughly equivalent to lateral 2.0-2.8 mm(37)), were processed for electron microscopy using osmium-thiocarbohydrazide-osmium post-fixation(38) (see **Supplement 1.4**). Samples were sectioned into ~70-75 nm ultrathin sections using an Ultracut UC7 ultramicrotome (Leica Biosystems). Three levels of section-rubans were collected at an interval of 10 μm, glued on a specimen mount with conductive carbon adhesive tabs (Electron Microscopy Sciences) and imaged by array tomography at 25 nm resolution with an acceleration voltage of 1.4 kV and current of 1.2 nA using a Zeiss Crossbeam 540 Gemini scanning EM (Zeiss) (3 images per embryo).

Images from the all 4 treatment groups were analyzed blind to the experimental conditions using QuPath (v0.2.0-m3) software (39). For each picture, region areas were traced and measured to calculate cell density. Cell type and apoptotic state was determined by ultrastructural features. Total cell numbers, dark cells (neuronal and glial cells), and apoptotic cells were then counted within the dorsal hippocampus (CA1, CA3, and dentate gyrus). The percentage of dark cell population or apoptotic cell number was calculated as a ratio over the total cell population (details in **Supplement 1.4**).

### 2.4 Statistical analyses

#### 2.4.1 Neuroimaging data analysis

Statistical analyses were performed using the R software package (R version 3.5.1, RMINC version 1.5.2.2 www.r-project.org). Once we confirmed there were no statistically significant differences between our two control groups, they were combined, leaving us with three treatment groups: saline (SAL), GD 9-poly I:C (POL E), and GD 17-poly I:C (POL L). To assess the effects of poly I:C exposure at different gestational timepoints on embryo neuroanatomy we ran a whole-brain voxel-wise linear mixed-effects model (‘mincLmer’; lme4_1.1-21 package; (40)) on the relative Jacobian determinant files using group and sex as fixed effects, and number of pups per litter, and cohort collection batch as random intercepts. A Satterthwaite approximation was used to compute degrees of freedom for every voxel (using the ‘mincLmerEstimateDF’ function). The False Discovery Rate (FDR) correction (using ‘mincFDR’) was applied to correct for multiple testing (41,42) (**Supplement 1.5.1** for details). This analysis was run again with the POL L group as the reference in order to directly compare POL E to POL L differences (**Supplement 2.2.1**). Sex differences were explored as a follow up analysis (**Supplement 1.5.1**). Putative differences in organ volume (i.e. lungs, heart, liver, etc) for the embryos were also investigated by applying deformation based morphometry (as described above) to the body cavity (**Supplement 1.5.1)**, however no differences were observed (**Supplement 2.2.2**).

#### 2.4.2 Electron microscopy

Given the small sample size and non-normal distribution of cell density measures collected, parametric statistics were not an appropriate choice to analyze the EM data (43). We chose to use a nonparametric Kruskall-Wallis test followed by a pairwise Wilcoxon test were used to assess group differences in the density of dark glial, dark neuronal, and apoptotic cells in the combined SAL, POL E, and POL L groups averaged per subject across slices (the SAL group was combined to maintain consistency with the MRI findings).

Given that the variance in cell density differed quite drastically between groups, we wanted to understand how the distributions in cell density differed between groups and how much, rather than a standard approach comparing differences in mean (standard t-test approach for normal distributions), or median (such as the Kruskal-Wallis test for non-normal distributions). In order to do so we applied the shift function to the density measures acquired from each slice per mouse (3) to maximize variance, rather than using the pooled data per mouse. The shift function (44) allows us to quantify how two distributions differ based on deciles of the distributions, i.e., it describes how one distribution should be transformed to match the other and estimates how and by how much one distribution must be shifted. When a significant difference is observed between deciles it suggests that there is a specific difference in the density of the cell type investigated (i.e., dark glia) between groups; this allows us to determine whether differences are consistent across the entire distribution, or more localized to one or both tails, or the center. In the context of the cell density data acquired, this technique allows us to compare groups beyond means or medians by accounting for the variance across the entire distribution of the data; this may provide us with a more nuanced understanding of differences between groups. Three pairwise comparisons were made (SAL – POL E, SAL – POL L, POL L – POL E) on the distributions for dark glia, dark neuron, and apoptotic cell density. A percentile bootstrap technique was used to derive confidence intervals based on differences in distribution at each decile of the distribution with a corresponding p-value. This was then repeated to assess sex differences, so the same comparisons were made in only males, and only females, followed by the same percentile bootstrap procedure.

## 3. Results

### 3.1 Embryo brain results

We observed a significant effect of GD9 MIA-exposure on the GD18 POL E embryo brain volume (t=4.242, <1%FDR), wherein POL E offspring had smaller volumes than SAL in the globus pallidus, hippocampus (including the dentate gyrus as well as more posterior regions), fornix, centromedian thalamic nucleus, and cerebellum. Larger volume was observed in the prelimbic area, lateral septum, subventricular zone, caudate-putamen, sexually dimorphic nucleus of the hypothalamus, basolateral amygdala, and CA1 region of the hippocampus in the POL E group relative to SAL (**Figure 2**).

**Figure 2.**
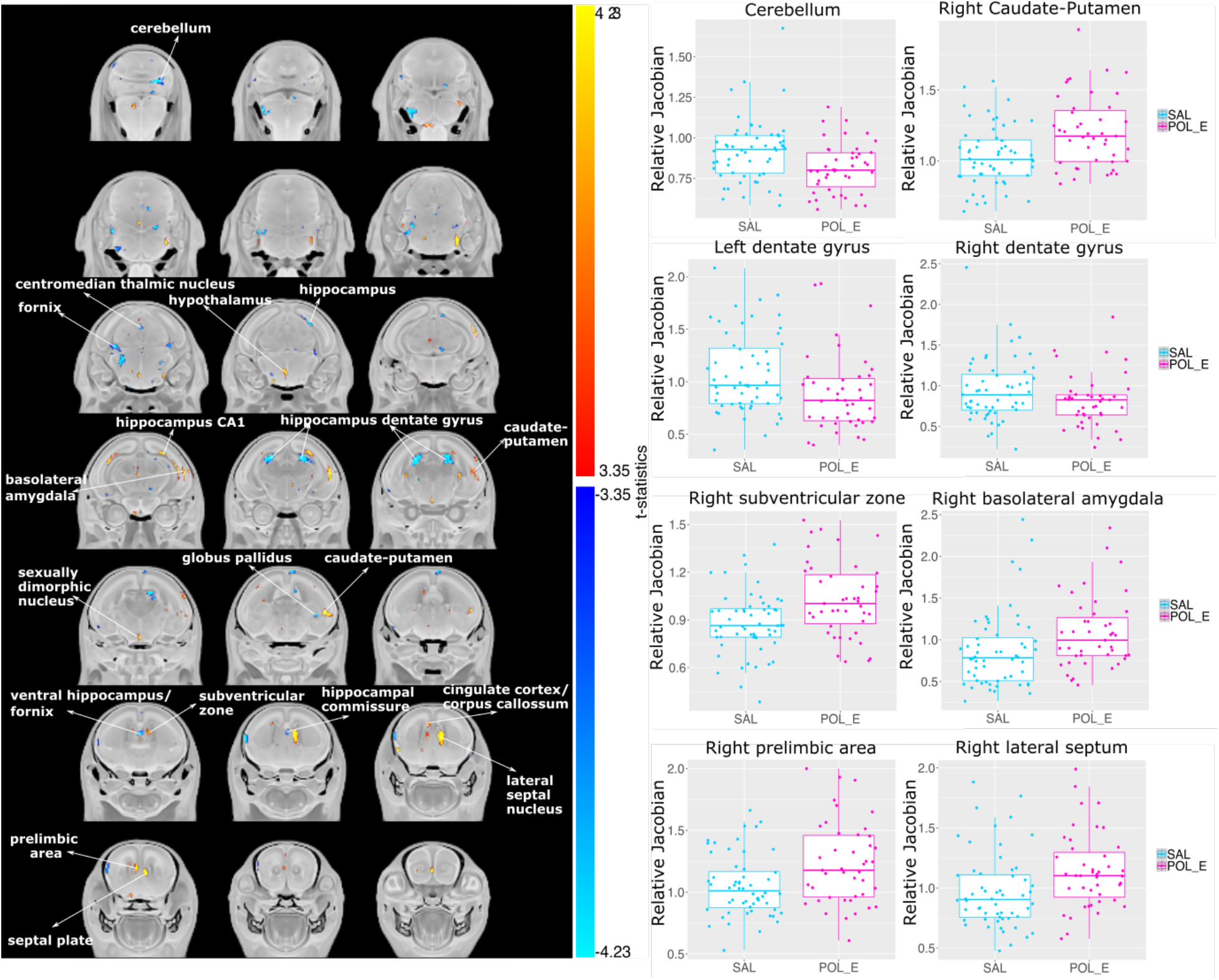
Neuroanatomical changes in the GD18 embryo brain following GD9-MIA exposure. **A.** t-statistic map of group (POL E vs SAL) thresholded at 5% (bottom, t=3.35) and 1% FDR (top, t=4.23) overlaid on the study average. **B.** Boxplots of peak voxels (voxels within a region of volume change showing largest effect) selected from regions of interest highlighted in white text in **A**. For all boxplots, the relative Jacobian determinants are plotted on the y-axis. Here a value of 1 means the voxel is no different than the average, anything above 1 is relatively larger, and below 1 is relatively smaller. For all boxplots, the midline represents the median of the data, the box represents the first and third quartiles, the vertical lines represent 1.5 x interquartile range of the data. Dots on the plot represent individual data points for each subject.

GD17 MIA-exposure induced very striking volumetric alterations, particularly volumetric increases in the brain of POL L offspring at GD18 (t=3.234, <1%FDR) relative to SAL offspring. Regions of volume increase include the ventral pallidum, septal plate and lateral septal nucleus, medial and lateral preoptic nuclei, caudate-putamen, globus pallidus, hippocampus (both dentate gyrus and CA1 regions), cingulum, anterior commissure, cortical plate, corpus callosum, external capsule, centromedian thalamus, and cerebellum. Decreases in volume were observed in the ventral hippocampus, more anterior subregions of the cortical plate, bilateral amygdala, fornix, and ventromedial thalamus (**Figure 3**).

**Figure 3.**
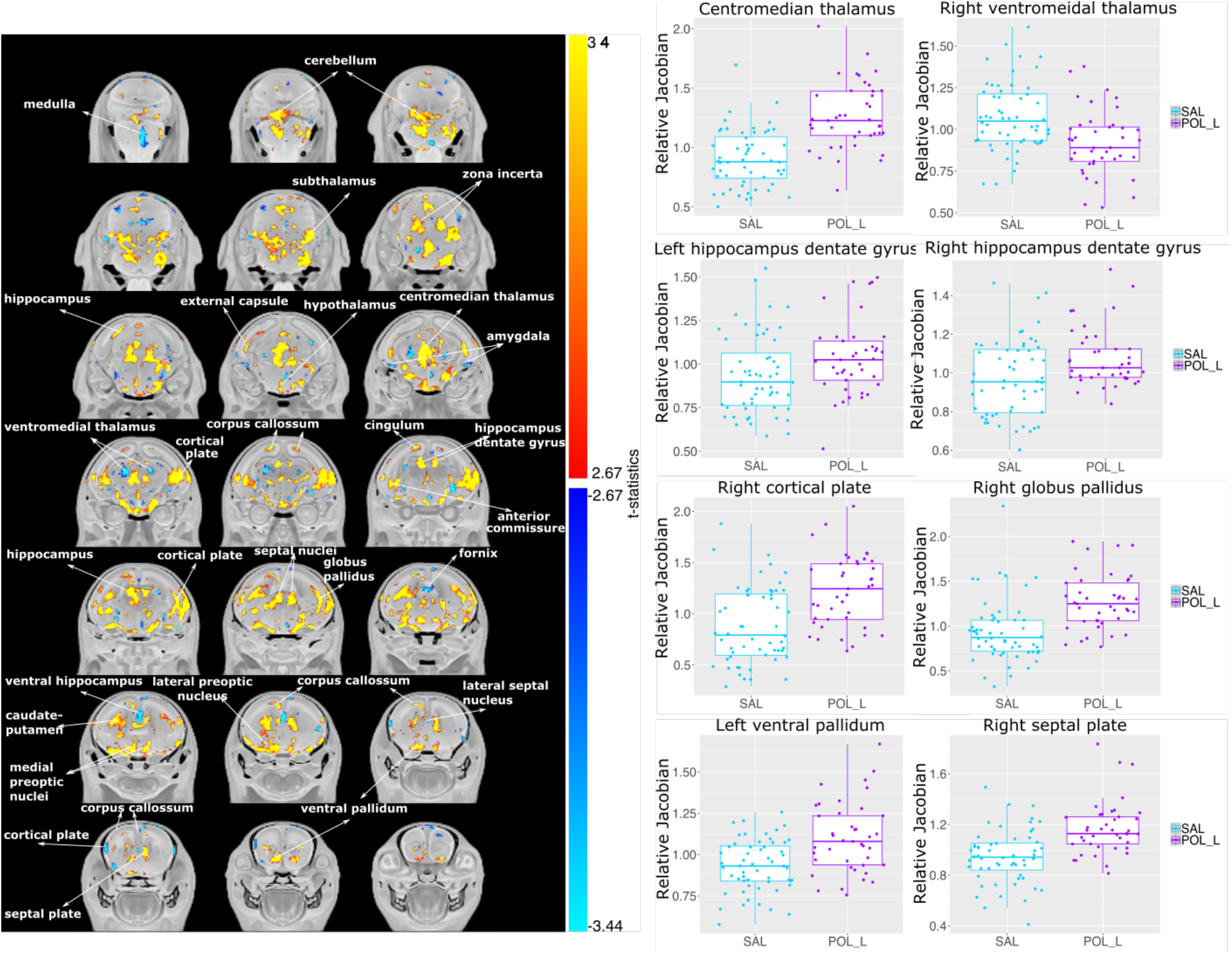
Neuroanatomical changes in the GD18 embryo brain following GD17-MIA exposure. **A.** t-statistic map of the group (POL L vs SAL) thresholded at 5% (bottom, t=2.67) and 1% FDR (top, t=3.44) overlaid on the study average. **B.** Boxplots of peak voxels (voxels within a region of volume change showing largest effect) selected from regions of interest highlighted in white text in **A**. For all boxplots, the relative Jacobian determinants are plotted on the y-axis as in Figure 2.

Interestingly, POL E and POL L MIA exposure were observed to have opposite effects on brain volume in some regions implicated in neurodevelopmental disorders and identified in previous MIA studies (11,14), such as the dorsal hippocampus, wherein GD9-exposure decreased volume, and GD17-exposure increased volume. Similar observations were made for the centromedian thalamic nucleus. The septal nucleus and caudate-putamen were increased in both MIA-exposed groups. A significant difference between POL E and POL L embryo brain anatomy was also observed (t=3.590, <1%FDR). These results are fully described in **Supplement 2.2.1** and **Supplementary Figure 1.** Post-hoc investigation of sex differences revealed no significant sex-by-group interactions.

### 3.2 Electron microscopy of embryo dorsal hippocampus

For aggregate density measure per group, there were no group differences in total cell density (chi-squared = 0.68038, df = 2, p-value = 0.7116; **Supplementary figure 3**). Although the density of dark glial and dark neuron cells appeared to be higher in the POL L offspring, there were no differences in dark glial cell density (chi-squared = 3.3221, df = 2, p-value = 0.1899), nor dark neuron cell density (chi-squared = 0.75759, df = 2, p-value = 0.6847). A significant group effect was observed for apoptotic cell density (chi-squared = 6.3491, df = 2, p-value = 0.04181) wherein the POL E offspring appeared to have greater density than the POL L offspring (p=0.053), as well as the SAL offspring, although not significantly so (p=0.117) (**Figure 4;** see **Supplementary figure 4** for more representative images of dark glia, dark neurons, and apoptotic cells).

**Figure 4.**
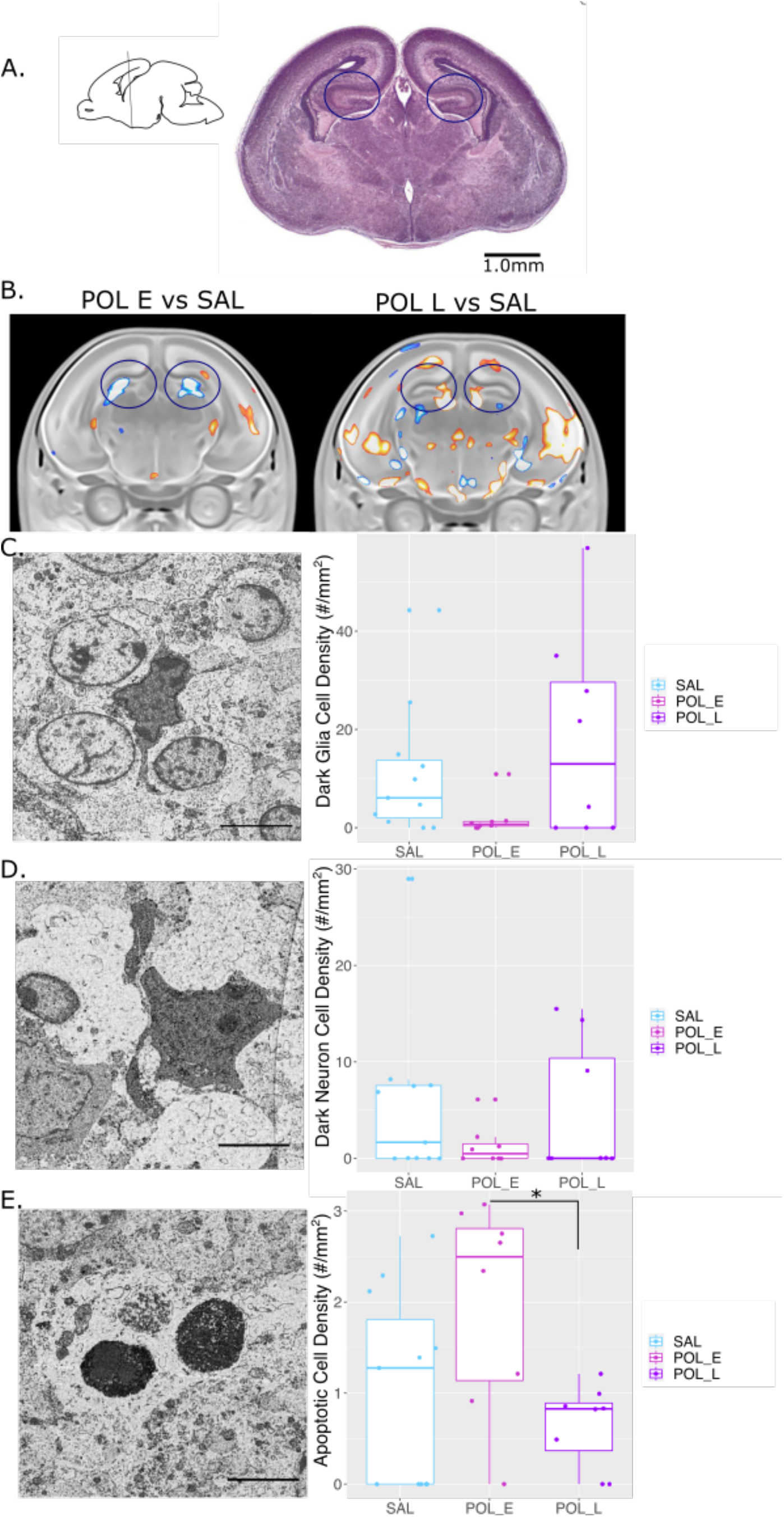
Differences in dark and apoptotic cell density with representative images captured using electron microscopy. **A.** Sagittal slice orienting to the region of the hippocampus selected, with the corresponding brain slice stained with Cresyl Violet from the GD18 mouse brain atlas (1mm scale bar), coronal slice 14 (36). The region of interest is highlighted in the circles. **B.** Representative slices of the hippocampus from the MRI results for POL E relative to SAL and POL L relative to SAL. The region of interest is highlighted in the circles. **C.** Image acquired by scanning electron microscopy (25 nm resolution) in the dorsal hippocampus from representative offspring (equivalent to coronal slice 14 from **A**) highlighting dark glial cells. Boxplot showing dark glial cell density (per mouse) per group (n=6-8/group). **D.** Representative dark neuron image with boxplot for dark neuron cell density per mouse. **E.** Representative apoptotic cell image with boxplot for apoptotic cell density per mouse. Scale bar equivalent to 5 μm. * p=0.053

Comparison of cell density distributions across groups confirmed that there were no overall differences in total cell density between groups (apart from a significant difference between POL E and SAL distributions only at the seventh decile of distribution (p=0.045) **Supplementary tables 2-4**).

Interestingly, dark glial density was significantly lower for POL E offspring relative to SAL at higher deciles of the distribution (5-9th decile, p<0.02; **Supplementary table 5; Figure 5A**), indicating that in general POL E offspring had decreased cell density. Distribution for POL L offspring were no different than SAL, however, they also had significantly more dark glia than POL E at higher deciles of the distribution (6-9th decile, p<0.04; **Supplementary tables 6 and 7**); in other words, POL L tend to have higher glial cell density than POL E, particularly among larger observations. Comparison of distributions for dark neurons revealed only subtle differences, with significantly higher density for POL E relative to SAL (decile 9, p=0.016; **Supplementary table 8**) and POL L (decile 8 and 9, p<0.01; **Supplementary table 10**) only at higher deciles of the distribution, with no differences between POL L and SAL (**Supplementary table 9; Figure 5B)**. Finally, for apoptotic cell density, POL E offspring had significantly higher density across lower deciles of distribution relative to SAL (deciles 1-4, p=0.039; **Supplementary table 11**), and across all deciles relative to POL L (p<0.02, **Supplementary table 12**); this indicates that among smaller observations, POL E tend to have higher apoptotic cell density than SAL, and higher density than POL L. No differences between POL L and SAL were observed (**Supplementary Table 13; Figure 5C**).

**Figure 5.**
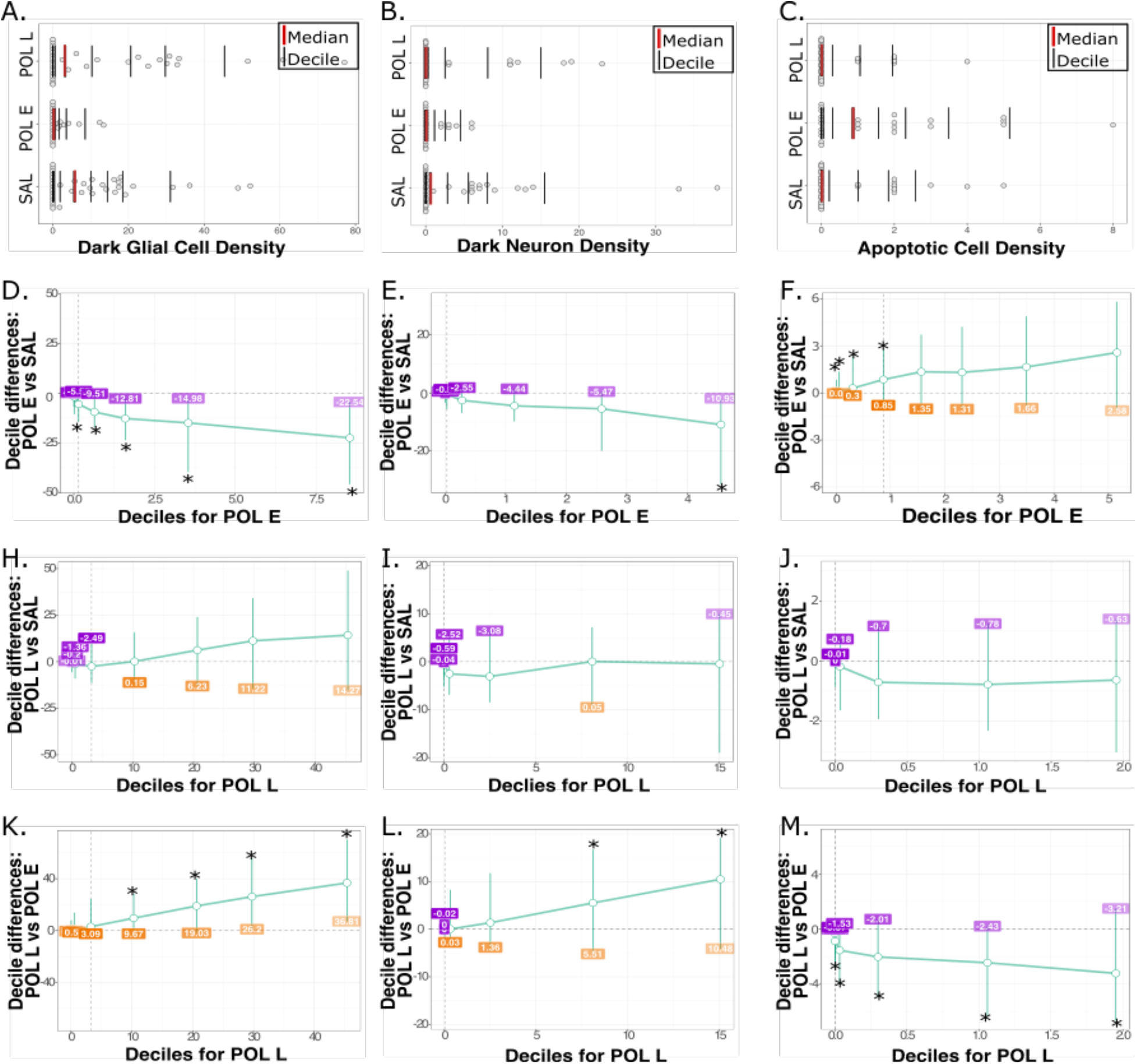
Differences in distribution of dark glial, dark neuron, and apoptotic cell density per group. Distribution of dark glial cell density **(A),** dark neuron density (**B**), and apoptotic cell density (**C**) for all hippocampal slices per animal. The red line identifies the median of the data, while each black bar denotes a decile of distribution. A percentile bootstrapping technique applied to identify the difference in decile between the POL E and SAL for dark glial cell density (**D**) dark neuron density (**E**), and apoptotic cell density (**F**) showing decreased density of dark glia, and increased density of apoptotic cells for POL E offspring. Next, POL L and SAL comparisons are shown for dark glia (**H**), dark neurons (**I**), and apoptotic cells (**J**). Finally, POL E vs POL L density is shown for dark glia (**K**), dark neurons (**L**), and apoptotic cells (**M**) showing that POL L had higher density that POL E for dark glia and neurons, while POL L has lower apoptotic cell density. *p<0.05

Sex differences in distribution were also observed for all cell types (total, dark glia, dark neurons, and apoptotic cells). Of interest, increased density of dark glial cells was observed in the POL L females relative to SAL, while decreased density was observed for the POL E females, further described in **Supplement 2.3.2** and **Supplementary Figure 5,** with the summary of results per decile provided in **Supplementary tables 14-37.**

## 4. Discussion

There is a well-established link between MIA-exposure *in utero* and latent neuroanatomical and behavioural abnormalities that emerge in adolescence or adulthood, with relevance to schizophrenia and ASD pathology (45–47). However, limited work has been conducted in the early neurodevelopmental period (12). We leveraged high-resolution *ex vivo* MRI and EM to characterize the effects of MIA-exposure at two gestational timepoints on the embryo brain at GD18. Our results suggest that the embryo mouse brain undergoes significant remodeling in response to MIA, particularly due to late gestational exposure, coupled with changes in the presence of dark and apoptotic cells in the hippocampus. Elucidating the neurodevelopmental changes across the embryonic periods following MIA-exposure is an important step towards our understanding of MIA-exposure as a primer downstream psychopathology and as a risk factor for an array of neuropsychiatric disorders.

Interestingly, we see volume reductions due to early MIA-exposure in a number of brain regions where we see striking volume expansions with late exposure. Since the late exposed embryo brains were harvested 24 hours following immune exposure, we are likely capturing an acute neuroinflammatory or stress response, or an acceleration of brain development in response to the immune stimulus. However, given that there was no difference based on SAL timing, it is likely that any acute effects are attributable to the MIA itself. Interestingly, there is homology between regions affected in the embryo brains, and those in which we observed altered neurodevelopmental trajectories from childhood to adulthood in our previously published work (11). Some of these regions include the striatum/caudate-putamen, hippocampus, lateral septum, cingulate cortex, and cerebellum, many of which have been implicated in neuroimaging studies of humans with schizophrenia or ASD (30,48,49). Previous animal studies also report that MIA in late gestation increases neuroinflammation in the embryo rat brain and decreases placental function as measured by T2-signal intensity (50). These findings suggest that increased neuroinflammation and decreased placental function could, in part, be driving some of the volumetric increases in the late exposed embryo brain, which may provide some mechanisms underlying disease pathology.

To gain more insight into the putative cellular underpinnings of the volumetric changes, we performed EM experiments in the dorsal hippocampus, a region highly implicated in neurodevelopmental pathology (31). In addition to the current findings, we previously observed interesting transcriptional changes to genes involved in early neurodevelopmental processes in the dorsal hippocampus of adolescent MIA-exposed mice – although following a different gestational exposure (11). Identifying the cellular processes triggered by MIA-exposure is critical to our understanding of how this risk factor may alter offspring neurodevelopmental trajectories. By leveraging high-resolution EM techniques, we have an unprecedented opportunity to investigate the brain parenchyma at nanoscale resolution (28). This allows for identification of changes in different cell types and their unique features. We identified differential effects due to gestational MIA-timing. In GD9-exposed offspring, where decreased dorsal hippocampal volume was detected, we observed an increase in apoptotic cell density and decrease in dark glial cell density compared to SAL offspring. Conversely, in GD17-exposed offspring, who had enlarged dorsal hippocampal volumes, there was a tendency for greater dark cell density (although not significantly), particularly for dark glial cells in the late exposed females. These cellular results align well with the volume-based MRI results wherein increased apoptosis could be linked to the decreased volume in GD9-exposed offspring, while increased dark cell density could be linked to an acute inflammatory response and increased volume in the GD17-exposed offspring. Importantly, this may point to differences in neuropathological mechanisms in the fetal brain associated with MIA-exposure, and to some putative sex differences that require further investigation.

We focused our analyses on dark cells, both neurons and glia, as well as apoptotic cells as these have been frequently detected in response to stress (29), aging (51), and neurodegeneration (52). They are thought to play a role in both normal and pathological synapse and neuronal network formation (52). Dark neurons are typically defined by a darker appearance, with an accumulation of mitochondria and nuclear indentations, associated with structural plasticity (52–54), as well as markers of cellular stress (dilated endoplasmic reticulum and Golgi apparatus) (28). Dark glial cells, particularly microglia, also display cellular markers of stress and have been shown to have hyper-ramified processes, which often leads to increased physiologically relevant contents such as synaptic contacts and increased phagocytic capacity(27,55). Further, reports of microglial reactivity and density in the brains of prenatally immune-challenged animals early in life are mixed, with observations of increased density and motility, as well as no differences (56). By focusing specifically on dark microglial cells in the future, we may gain better insight into the phenotypic variability of these reactive cells, and parse some of the heterogeneity in the literature.

At GD9, corresponding with our first MIA-exposure, microglia colonize the brain, initiating their development towards maturity. Interestingly, sex differences in the number and morphology of microglia have been observed with males showing greater numbers of these cells earlier in development (PND4) than females, who have more microglia later in development (PND30-60) (57). GD9 also occurs at a time at which the fetal brain is undergoing extensive neural proliferation and migration, which transitions more towards circuit refinement and cortical organization by GD17 (our late MIA-timepoint) (58). Importantly, microglia may play important roles in the regulation of apoptosis (59) as well as the permeability and formation of the blood-brain barrier, which typically takes shape between GD13.5-15.5 in the mouse. Apoptosis is a critical cellular process in early brain and placental development *in utero* (60). The process of apoptosis has been detected as early as GD5, but increases significantly towards the end of gestation, peaking in early postnatal life before dropping off (61). In contrast with our findings, previous rodent studies using immunohistochemical techniques have actually observed an increase in apoptosis following lipopolysaccharide exposure in late pregnancy in rats (62), and poly I:C exposure at GD17, but not GD9 in mice (22). This discrepancy between these findings and ours may be due to differences in techniques or features used to identify these cells; immunohistochemistry may be less sensitive to detection of ongoing apoptosis.

The results presented here should be considered in light of their limitations. The design of our embryo study would be more complete with an assessment of neuroanatomy acutely following the GD9-exposure at GD10; this would allow us to detect whether volume increases, as those detected in the GD17-exposed offspring, are a response to acute inflammation, or specific to that gestational time point. Additionally, examining the brain of GD17-exposed offspring 9 days following MIA exposure, at PND8, could provide a similar delay after MIA before collection for imaging, however, the comparison of embryonic vs. postnatal brain development may have its own confounds. Unfortunately, the embryo is too small for MRI acquisition at GD 9 or 10, however, imaging could be performed using other techniques such as optical projection tomography (63). Regarding the EM analysis, we did not look at the extracellular space, which could contribute to the changes in volume detected in the MRI. In future work, cryofixation methods for preserving the EM samples would allow for more in depth analyses of the extracellular space volume and composition (64).

We comprehensively examined the effects of prenatal MIA-exposure, a known risk factor for neuropsychiatric disorders, at two gestational timepoints on embryo brain anatomy at the gross morphological and cellular levels. We identified striking neuroanatomical remodeling in the embryo brain, particularly following exposure in late gestation; we also observed sex-dependent alterations in the density of dark neuronal and glial cells in the dorsal hippocampus, with greater cell density in female offspring. This may reflect the initiation of pathological circuit remodeling. These findings show that MIA-exposure induces striking neurodevelopmental changes in embryonic development, which may further our understanding of how this risk factor increases the likelihood of developing neuropsychiatric illnesses later in life.

## Supporting information

Supplementary material, results, figures and tables

## Acknowledgements

The authors are grateful to Roulin Gao for providing training in how to harvest embryos. We would like to thank Dr’s Bruno Giros and Salah El Mestikawy for lending us their centrifuge. Finally, the authors would like to acknowledge their funding bodies, including the Canadian Institute of Health Research and Healthy Brains for Healthy Lives for providing support for this research. Additionally, we would like to thank the Fonds de Recherche du Québec en Santé for providing salary support for EG, KP, and MMC, as well as the Kappa Kappa Gamma Foundation of Canada for supporting EG’s salary. MET is a Tier 2 Canada Research Chair in *Neurobiology of Aging and Cognition*.

## References

1. Brown AS, Cohen P, Harkavy-Friedman J, Babulas V, Malaspina D, Gorman JM, et al. Prenatal rubella, premorbid abnormalities, and adult schizophrenia. Biol Psychiatry. 2001 Mar 15;49(6):473–86.

2. Brown AS, Begg MD, Gravenstein S, Schaefer CA, Wyatt RJ, Bresnahan M, et al. Serologic evidence of prenatal influenza in the etiology of schizophrenia. Arch Gen Psychiatry. 2004 Aug;61(8):774–80.

3. Selten JP, van Duursen R, van der Graaf Y, Gispen-de Wied C, Kahn RS. 736 – Second-trimester exposure to maternal stress is a possible risk factor for psychotic illness in the child. Schizophr Res. 1997 Jan 1;24(1):258.

4. Canetta S, Brown A. Prenatal infection, maternal immune activation, and risk for schizophrenia [Internet]. Vol. 3, Translational Neuroscience. 2012. Available from: http://dx.doi.org/10.2478/s13380-012-0045-6

5. Choi GB, Yim YS, Wong H, Kim S, Kim H, Kim SV, et al. The maternal interleukin-17a pathway in mice promotes autism-like phenotypes in offspring [Internet]. Vol. 351, Science. 2016. p. 933–9. Available from: http://dx.doi.org/10.1126/science.aad0314

6. Gumusoglu SB, Stevens HE. Maternal Inflammation and Neurodevelopmental Programming: A Review of Preclinical Outcomes and Implications for Translational Psychiatry. Biol Psychiatry. 2019 Jan 15;85(2):107–21.

7. Solek CM, Farooqi N, Verly M, Lim TK, Ruthazer ES. Maternal immune activation in neurodevelopmental disorders [Internet]. Vol. 247, Developmental Dynamics. 2018. p. 588–619. Available from: http://dx.doi.org/10.1002/dvdy.24612

8. Boksa P. Effects of prenatal infection on brain development and behavior: a review of findings from animal models. Brain Behav Immun. 2010 Aug;24(6):881–97.

9. Thion MS, Ginhoux F, Garel S. Microglia and early brain development: An intimate journey. Science. 2018 Oct 12;362(6411):185–9.

10. Reisinger S, Khan D, Kong E, Berger A, Pollak A, Pollak DD. The Poly(I:C)-induced maternal immune activation model in preclinical neuropsychiatric drug discovery. Pharmacol Ther. 2015 May 1;149:213–26.

11. Guma E, Bordignon P do C, Devenyi GA, Gallino D, Anastassiadis C, Cvetkovska V, et al. Early or late gestational exposure to maternal immune activation alters neurodevelopmental trajectories in mice: an integrated neuroimaging, behavioural, and transcriptional study. Biol Psychiatry [Internet]. 2021 Mar 23; Available from: https://www.sciencedirect.com/science/article/pii/S0006322321011768

12. Guma E, Plitman E, Chakravarty MM. The role of maternal immune activation in altering the neurodevelopmental trajectories of offspring: A translational review of neuroimaging studies with implications for autism spectrum disorder and schizophrenia. Neurosci Biobehav Rev [Internet]. 2019; Available from: https://www.sciencedirect.com/science/article/pii/S0149763419302088

13. Ellman LM, Deicken RF, Vinogradov S, Kremen WS, Poole JH, Kern DM, et al. Structural brain alterations in schizophrenia following fetal exposure to the inflammatory cytokine interleukin-8. Schizophr Res. 2010 Aug;121(1-3):46–54.

14. Crum WR, Sawiak SJ, Chege W, Cooper JD, Williams SCR, Vernon AC. Evolution of structural abnormalities in the rat brain following in utero exposure to maternal immune activation: A longitudinal in vivo MRI study. Brain Behav Immun. 2017 Jul;63:50–9.

15. Piontkewitz Y, Arad M, Weiner I. Abnormal trajectories of neurodevelopment and behavior following in utero insult in the rat. Biol Psychiatry. 2011 Nov 1;70(9):842–51.

16. Graham AM, Rasmussen JM, Rudolph MD, Heim CM, Gilmore JH, Styner M, et al. Maternal Systemic Interleukin-6 During Pregnancy Is Associated With Newborn Amygdala Phenotypes and Subsequent Behavior at 2 Years of Age. Biol Psychiatry. 2018 Jan 15;83(2):109–19.

17. Rudolph MD, Graham AM, Feczko E, Miranda-Dominguez O, Rasmussen JM, Nardos R, et al. Maternal IL-6 during pregnancy can be estimated from newborn brain connectivity and predicts future working memory in offspring. Nat Neurosci. 2018 May;21(5):765–72.

18. Spann MN, Monk C, Scheinost D, Peterson BS. Maternal Immune Activation During the Third Trimester Is Associated with Neonatal Functional Connectivity of the Salience Network and Fetal to Toddler Behavior. J Neurosci. 2018 Mar 14;38(11):2877–86.

19. Canales CP, Estes ML, Cichewicz K, Angara K, Aboubechara JP, Cameron S, et al. Sequential perturbations to mouse corticogenesis following in utero maternal immune activation. Elife [Internet]. 2021 Mar 5;10. Available from: http://dx.doi.org/10.7554/eLife.60100

20. Lombardo MV, Moon HM, Su J, Palmer TD, Courchesne E, Pramparo T. Maternal immune activation dysregulation of the fetal brain transcriptome and relevance to the pathophysiology of autism spectrum disorder. Mol Psychiatry. 2018 Apr;23(4):1001–13.

21. Kalish BT, Kim E, Finander B, Duffy EE, Kim H, Gilman CK, et al. Maternal immune activation in mice disrupts proteostasis in the fetal brain. Nat Neurosci. 2021 Feb;24(2):204–13.

22. Meyer U, Nyffeler M, Engler A, Urwyler A, Schedlowski M, Knuesel I, et al. The time of prenatal immune challenge determines the specificity of inflammation-mediated brain and behavioral pathology. J Neurosci. 2006 May 3;26(18):4752–62.

23. Cavalcante MB, Cavalcante CT de MB, Sarno M, Barini R, Kwak-Kim J. Maternal immune responses and obstetrical outcomes of pregnant women with COVID-19 and possible health risks of offspring. J Reprod Immunol. 2021 Feb;143:103250.

24. Reyes-Lagos JJ, Abarca-Castro EA, Echeverría JC, Mendieta-Zerón H, Vargas-Caraveo A, Pacheco-López G. A Translational Perspective of Maternal Immune Activation by SARS-CoV-2 on the Potential Prenatal Origin of Neurodevelopmental Disorders: The Role of the Cholinergic Anti-inflammatory Pathway [Internet]. Vol. 12, Frontiers in Psychology. 2021. Available from: http://dx.doi.org/10.3389/fpsyg.2021.614451

25. Wu D, Zhang J. Recent Progress in Magnetic Resonance Imaging of the Embryonic and Neonatal Mouse Brain. Front Neuroanat. 2016 Mar 3;10:18.

26. Barron HC, Mars RB, Dupret D, Lerch JP, Sampaio-Baptista C. Cross-species neuroscience: closing the explanatory gap. Philos Trans R Soc Lond B Biol Sci. 2021 Jan 4;376(1815):20190633.

27. Bisht K, Sharma KP, Lecours C, Sánchez MG, El Hajj H, Milior G, et al. Dark microglia: A new phenotype predominantly associated with pathological states. Glia. 2016 May;64(5):826–39.

28. Nahirney PC, Tremblay M-E. Brain Ultrastructure: Putting the Pieces Together. Front Cell Dev Biol. 2021 Feb 18;9:629503.

29. Henry MS, Bisht K, Vernoux N, Gendron L, Torres-Berrio A, Drolet G, et al. Delta Opioid Receptor Signaling Promotes Resilience to Stress Under the Repeated Social Defeat Paradigm in Mice. Front Mol Neurosci. 2018 Apr 6;11:100.

30. van Erp TGM, Hibar DP, Rasmussen JM, Glahn DC, Pearlson GD, Andreassen OA, et al. Subcortical brain volume abnormalities in 2028 individuals with schizophrenia and 2540 healthy controls via the ENIGMA consortium. Mol Psychiatry. 2016 Apr;21(4):547–53.

31. Lieberman JA, Girgis RR, Brucato G, Moore H, Provenzano F, Kegeles L, et al. Hippocampal dysfunction in the pathophysiology of schizophrenia: a selective review and hypothesis for early detection and intervention. Mol Psychiatry. 2018 Aug;23(8):1764–72.

32. Hui CW, St-Pierre A, El Hajj H, Remy Y, Hébert SS, Luheshi GN, et al. Prenatal Immune Challenge in Mice Leads to Partly Sex-Dependent Behavioral, Microglial, and Molecular Abnormalities Associated with Schizophrenia. Front Mol Neurosci. 2018;11:13.

33. Spencer Noakes TL, Henkelman RM, Nieman BJ. Partitioning k-space for cylindrical three-dimensional rapid acquisition with relaxation enhancement imaging in the mouse brain. NMR Biomed [Internet]. 2017 Nov;30(11). Available from: http://dx.doi.org/10.1002/nbm.3802

34. Avants BB, Tustison NJ, Song G, Cook PA, Klein A, Gee JC. A reproducible evaluation of ANTs similarity metric performance in brain image registration. Neuroimage. 2011 Feb 1;54(3):2033–44.

35. Chung MK, Worsley KJ, Paus T, Cherif C, Collins DL, Giedd JN, et al. A unified statistical approach to deformation-based morphometry. Neuroimage. 2001 Sep;14(3):595–606.

36. Schambra U. Prenatal Mouse Brain Atlas: Color images and annotated diagrams of: Gestational Days 12, 14, 16 and 18 Sagittal, coronal and horizontal section. Springer, Boston, MA; 2008.

37. Franklin KB, Paxinos G. The Mouse Brain in Stereotaxic Coordinates, Compact. The Coronal Plates and Diagrams. Amsterdam: Elsevier Academic Press; 2008.

38. Ellisman M, Shu X, Lev-Ram V, Deerinck T, Tsien R, Lamont S, et al. Bridging Gaps in Imaging by Applying EM Tomography and Serial Block Face SEM, Including a New Genetically Encoded Tag for Correlated Light and 3D Electron Microscopy of Intact Cells, Tissues and Organisms: Integrating the Resulting Correlated Image Data Using the Whole Brain Catalog [Internet]. Vol. 17, Microscopy and Microanalysis. 2011. p. 2–3. Available from: http://dx.doi.org/10.1017/s1431927611000882

39. Bankhead P, Loughrey MB, Fernández JA, Dombrowski Y, McArt DG, Dunne PD, et al. QuPath: Open source software for digital pathology image analysis. Sci Rep. 2017 Dec 4;7(1):16878.

40. Bates D, Mächler M, Bolker B, Walker S. Fitting Linear Mixed-Effects Models Usinglme4 [Internet]. Vol. 67, Journal of Statistical Software. 2015. Available from: http://dx.doi.org/10.18637/jss.v067.i01

41. Benjamini Y, Hochberg Y. Controlling the False Discovery Rate: A Practical and Powerful Approach to Multiple Testing. J R Stat Soc Series B Stat Methodol. 1995;57(1):289–300.

42. Genovese CR, Lazar NA, Nichols T. Thresholding of statistical maps in functional neuroimaging using the false discovery rate. Neuroimage. 2002 Apr;15(4):870–8.

43. Fagerland MW. t-tests, non-parametric tests, and large studies--a paradox of statistical practice? BMC Med Res Methodol. 2012 Jun 14;12:78.

44. Rousselet GA, Pernet CR, Wilcox RR. Beyond differences in means: robust graphical methods to compare two groups in neuroscience. Eur J Neurosci. 2017 Jul;46(2):1738–48.

45. Knuesel I, Chicha L, Britschgi M, Schobel SA, Bodmer M, Hellings JA, et al. Maternal immune activation and abnormal brain development across CNS disorders. Nat Rev Neurol. 2014 Nov;10(11):643–60.

46. Estes ML, Kimberley McAllister A. Maternal immune activation: Implications for neuropsychiatric disorders. Science. 2016 Aug 19;353(6301):772–7.

47. Brown AS, Meyer U. Maternal Immune Activation and Neuropsychiatric Illness: A Translational Research Perspective. Am J Psychiatry. 2018 Nov 1;175(11):1073–83.

48. van Erp TGM, Walton E, Hibar DP, Schmaal L, Jiang W, Glahn DC, et al. Cortical Brain Abnormalities in 4474 Individuals With Schizophrenia and 5098 Control Subjects via the Enhancing Neuro Imaging Genetics Through Meta Analysis (ENIGMA) Consortium. Biol Psychiatry. 2018 Nov 1;84(9):644–54.

49. van Rooij D, Anagnostou E, Arango C, Auzias G, Behrmann M, Busatto GF, et al. Cortical and Subcortical Brain Morphometry Differences Between Patients With Autism Spectrum Disorder and Healthy Individuals Across the Lifespan: Results From the ENIGMA ASD Working Group. Am J Psychiatry. 2018 Apr 1;175(4):359–69.

50. Girard S, Tremblay L, Lepage M, Sébire G. IL-1 receptor antagonist protects against placental and neurodevelopmental defects induced by maternal inflammation. J Immunol. 2010 Apr 1;184(7):3997–4005.

51. Tremblay M-È, Zettel ML, Ison JR, Allen PD, Majewska AK. Effects of aging and sensory loss on glial cells in mouse visual and auditory cortices. Glia. 2012 Apr;60(4):541–58.

52. Bisht K, Sharma K, Lacoste B, Tremblay M-È. Dark microglia: Why are they dark? Commun Integr Biol. 2016 Oct 3;9(6):e1230575.

53. Versaevel M, Braquenier J-B, Riaz M, Grevesse T, Lantoine J, Gabriele S. Super-resolution microscopy reveals LINC complex recruitment at nuclear indentation sites. Sci Rep. 2014 Dec 8;4:7362.

54. Zimatkin SM, Bon’ EI. Dark Neurons of the Brain. Neurosci Behav Physiol. 2018 Oct 1;48(8):908–12.

55. St-Pierre M-K, Šimončičová E, Bögi E, Tremblay M-È. Shedding Light on the Dark Side of the Microglia. ASN Neuro. 2020 Jan;12:1759091420925335.

56. Smolders S, Notter T, Smolders SMT, Rigo J-M, Brône B. Controversies and prospects about microglia in maternal immune activation models for neurodevelopmental disorders. Brain Behav Immun. 2018 Oct;73:51–65.

57. Schwarz JM, Sholar PW, Bilbo SD. Sex differences in microglial colonization of the developing rat brain. J Neurochem. 2012 Mar;120(6):948–63.

58. Selemon LD, Zecevic N. Schizophrenia: a tale of two critical periods for prefrontal cortical development. Transl Psychiatry. 2015 Aug 18;5:e623.

59. Ozaki K, Kato D, Ikegami A, Hashimoto A, Sugio S, Guo Z, et al. Maternal immune activation induces sustained changes in fetal microglia motility. Sci Rep. 2020 Dec 7;10(1):21378.

60. Mor G, Abrahams VM. Potential role of macrophages as immunoregulators of pregnancy. Reprod Biol Endocrinol. 2003 Dec 2;1:119.

61. Mosley M, Shah C, Morse KA, Miloro SA, Holmes MM, Ahern TH, et al. Patterns of cell death in the perinatal mouse forebrain. J Comp Neurol. 2017 Jan 1;525(1):47–64.

62. Cai Z, Pan ZL, Pang Y, Evans OB, Rhodes PG. Cytokine induction in fetal rat brains and brain injury in neonatal rats after maternal lipopolysaccharide administration. Pediatr Res. 2000 Jan;47(1):64–72.

63. Sharpe J. Optical projection tomography as a new tool for studying embryo anatomy. J Anat. 2003 Feb;202(2):175–81.

64. Korogod N, Petersen CCH, Knott GW. Ultrastructural analysis of adult mouse neocortex comparing aldehyde perfusion with cryo fixation. Elife [Internet]. 2015 Aug 11;4. Available from: http://dx.doi.org/10.7554/eLife.05793

