## Supplementary material, results, figures and tables for "Differential effects of early or late exposure to prenatal maternal immune activation on mouse embryonic neurodevelopment"

### Supplementary Materials

#### 1. Supplementary methods

##### 1.1 Animals and maternal immune activation (MIA)

C57BL/6J mice were used throughout the study and were bred in our facility under a 12-hour light cycle (8am-8pm), with food and water access *ad libitum*. C57BL/6J female and male mice of breeding age (8-12 weeks old) were subject to timed mating procedures. One male and one female were placed in a new cage in the afternoon, checked for the appearance of a plug, weighed, and separated the following morning. When a seminal plug was observed, this was considered gestational day (GD) 0 (mice were allowed only 1 night together to improve the accuracy of GD 0 detection).

For embryo sample collection, pregnant dams were randomly assigned to one of four treatment groups: (1) poly I:C (P1530-25MG polyinosinic:polycytidylic acid sodium salt TLR ligand tested; Sigma Aldrich) (5mg/kg, intraperitoneally) at gestational day (GD) 9 (POL E; n = 7 dams [4 cohort 1/ 3 cohort 2]), (2) 0.9% sterile NaCl solution at GD 9 (SAL E; n = 7 dams [6 cohort 1/ 1 cohort 2]), (3) poly I:C at GD17 (POL L; n = 7 dams [6 cohort 1/ 2 cohort 2], or (4) saline at GD17 (SAL L; n = 4 dams [2 cohort 1/ 2 cohort 2]). Collection was performed in two rounds (cohorts) with two different batches of poly I:C (same product). **Table 1** outlines total sample size for embryo groups, with those from cohort 2 in brackets.

In a separate group of dams, poly I:C or saline was injected as described above (GD 9-POL: n=3 batch 1, n=4 batch 2; GD 17-POL n=3 batch 1, n=4 batch 2; GD 9-SAL n=5, GD 17-SAL n=3). Three hours following injection, dams were sacrificed by decapitation without anesthesia, and trunk blood was collected in a 1.5 mL Eppendorf tube. The blood was allowed to coagulate at room temperature for 30 minutes, and then centrifuged for 10 minutes at 4 ℃, with 2000 revolutions per minute. Serum was collected and stored at -80 ℃ until ready for analysis. Serum samples were shipped to the University of Maryland Core Cytokine Facility (http://www.cytokines.com/) for multiplex ELISA to measure levels of IL-6, TNF-α, IL-1β, IL-10 in order to assess the immunostimulatory potential of our poly I:C model. We chose to use a separate group of dams to ensure we could collect enough blood for analysis, and so as not to introduce an additional stressful experience for the dam, thereby potentially confounding the neurodevelopmental trajectory of offspring. Detection ranges were as follows IL-6 (0.64-8000 pg/ml), TNF-α (0.64-3500 pg/ml), IL-1β (0.64-15000 pg/ml), IL-10 (0.64-20000 pg/ml).

##### 1.2 Brain sample preparation

Embryos were harvested at GD 18 as follows. Pregnant dams were euthanized by cervical dislocation without anesthesia, embryos were removed and placed in ice cold 1x phosphate buffered saline (PBS) solution. One at a time, embryos were moved into warm (37 °C) PBS, separated from the uterus and placenta, and allowed to bleed out under agitation. A piece of the yolk sac was collected for genotyping for each embryo in order to identify the sex of the mouse via presence of the SRY gene (performed by Transnetyx, Memphis, TN). Once blood was successfully drained, embryos were placed in 4% paraformaldehyde (PFA) with 2% gadolinium (MRI contrast agent; Bracco Imaging S.p.A) in PBS, and post-fixed for 1 week at 4 °C. Finally, embryos were moved to a 2% gadolinium, 0.02% sodium azide 1x PBS solution for long-term storage at 4 °C until scanning.

##### 1.3 Magnetic resonance image

###### 1.3.1 Acquisition

Prior to imaging, the samples were removed from the contrast agent solution, blotted and placed in 13-mm diameter plastic tubes filled with a proton-free susceptibility-matching fluid (Fluorinert FC-77, 3M Corp. St. Paul, MN). An anatomical scan was performed using a T2-weighted, 3D fast spin echo sequence using a cylindrical k-space acquisition (Nieman, et al., 2005) with TR/TE=350/12 ms, echo train length=6, two averages, field-of-view 20 mm x 20 mm x 25 mm, matrix size=504 x 504 x 630 (Spencer Noakes, Henkelman, and Nieman 2017). Total imaging time was 14 hours producing images with 40 μm isotropic resolution. MR images of precision-machined phantoms were aligned towards a computed tomography (CT) scan of the same phantom to produce distortion correcting transformations to correct for geometric distortions in a coil-specific manner.

###### 1.3.2 MRI processing

Images were first cropped to center around the head of the embryos, an N4 correction for B1 bias field inhomogeneities (Tustison et al. 2010) and denoising using non-local means (minc_anlm) (Manjón et al. 2010) was applied, and the background was set to zero using minc tools. Next, brain images of all subjects in the study were aligned using the antsMultivariateTemplateConstruction2.sh tool (https://github.com/CoBrALab/twolevel_ants_dbm) (Avants et al. 2011). Briefly, images were aligned using rigid registration (translation and rotation), followed by affine (rigid, scaling, and shear), and finally, by nonlinear registration yielding a precise anatomical alignment in an automated, minimally biased fashion, similar to what is done in adult mouse brain studies (Guma et al. 2021). The output of this iterative registration procedure is a study-specific average against which groups can be compared, as well as deformation fields that map each individual subject to the average at the voxel level. Further, the Jacobian determinants of each deformation field provide a measure of volume difference at each voxel in the image relative to the average (Chung et al. 2003). Relative Jacobian determinants (used for statistical analysis in this work) explicitly model only the non-linear part of the deformations and remove residual global linear transformations (attributable to differences in total brain size). Prior to performing statistics, Jacobian determinants were blurred at 0.16 mm full-width-at-half-maximum to better conform to Gaussian assumptions for downstream statistical testing.

##### 1.4 Electron microscopy

After post-fixation (described in the main text, **2.3**), tissues were incubated in a mixture (1:1) of 3% ferrocyanide (BioShop, cat# PFC232.250) combined with 4% osmium tetroxide (Electron Microscopy Sciences, cat#19170) for 1 hour, 1% thiocarbohydrazide (in PBS; Electron Microscopy Sciences, cat# 2231-57-4) for 20 minutes, and in 2% osmium tetroxide for 30 minutes. Osmium-thiocarbohydrazide-osmium post-fixed sections were then dehydrated (ethanol (2× in 35%, 1× in 50%, 1× in 70%, 1× in 80%, 1× in 90%, 2× in 100%) followed by 3× in propylene oxide, for 5 minutes each), infiltrated with Durcupan ACM resin (MilliporeSigma, cat# 44611-44614) for 24 hours, and embedded between two fluoropolymer ACLAR® sheets (Electron Microscopy Sciences, cat# 50425-25) at 55-60 °C for 5 days. Using a binocular microscope, the dorsal hippocampus was excised from tissue sections using a razor blade, glued with Ultra Gel Control super glue (base of ethylene cyanoacrylate, LePage) onto a resin block, and left to incubate overnight at 55-60 °C. Next, the resin block was trimmed with a razor blade into a pyramidal shape to allow for more stable and precise sectioning. Following glue removal, tissue was sectioned into ~70-75 nm ultrathin sections using an Ultracut UC7 ultramicrotome (Leica Biosystems). Three levels of section-rubans were collected at an interval of 10 µm, which represented different levels of the dorsal hippocampus. Section-rubans were collected on a silicon nitride chip and glued on a specimen mounts and imaged by array tomography at 25 nm resolution with an acceleration voltage of 1.4 kV and current of 1.2 nA using a Crossbeam 540 Gemini scanning electron microscope (Zeiss).

Images from the all treatment groups were analyzed blind to the experimental conditions using QuPath software (v0.2.0-m3) (Bankhead et al. 2017). For each picture, region areas were traced using the "polygon" tool and measured to determine cell density. Total cell numbers (identified by a well-defined nucleus), dark cells (neuronal and glial cells), and apoptotic cells were then counted within the dorsal hippocampus (CA1, CA3, and dentate gyrus). Data were reported both as a measure of cell density (divided by the region area) or of cell population percentage (divided by the total cell count).

Dark cells were identified by their darker cytoplasm and dark nucleus with a faint heterochromatine pattern (Bisht et al. 2016; Zimatkin and Bon' 2018; St-Pierre et al. 2020). Dark neurons were distinguished by their often enlarged cell cytoplasm with a visible apical dendrite (St-Pierre et al. 2020) (see http://www.bu.edu/agingbrain/chapter-1-neuronal-cell-bodies-2 for more detail) . Of note, some dark neurons were, however, shrunked, extended and deformed but recognised by their pyknotic nuclei (Zimatkin and Bon' 2018). Glial cell bodies were often smaller in size than dark neurons as well as less regular with, occasionally, several smaller irregular protrusions. Moreover, in contrast to dark neurons, dark glial cell nuclei possess a distinct faint heterochromatin pattern, showing a thin rim of more condensed chromatin at the edge of the neuropil or clumped patches of chromatin (Peters, Palay, and Webster 1991). Lastly, apoptotic cells or dying cells were identified by a pyknotic or fragmented nucleus, a distended or incomplete nuclear envelope and an accumulation of autophagic vacuoles in their cytoplasm (Taatjes et al. 2008; Peters, Palay, and Webster 1991).

##### 1.5 Statistical analyses

###### 1.5.1 Neuroimaging analysis

Our first goal was to determine whether there were neuroanatomical differences in our two control groups, SAL E (GD9) and SAL L (GD17). We ran a whole-brain (delimited by a brain mask) voxel-wise linear mixed effects model on the relative Jacobian determinant files of each subject with injection timing and sex as fixed effects, and litter and number of pups per litter as random intercept to account for litter-specific variation. A False Discovery Rate (FDR) correction was applied. There were no differences, and SAL E and SAL L offspring were combined and set as the reference group. Finally, sex differences were explored, investigating the interaction between group and sex, with the same covariates as above. The statistical models applied to the relative Jacobian determinants to assess for group differences, described in **X** are detailed below:

**Main model:** $Y_{subject,j} = \beta_0 + \beta_1 sexF_{subject,j} + \beta_2 groupPOL\_E_{subject,j} + \beta_3 groupPOL\_L_{subject,j} + \mathbf{b_1} cohort + \mathbf{b_2} litter\ size + \epsilon_{subject,j}$

**Sex interaction model:** $Y_{subject,j} = \beta_0 + \beta_1 sexF_{subject,j} + \beta_2 groupPOL\_E_{subject,j} + \beta_3 groupPOL\_L_{subject,j} + \beta_4 sexF{:}groupPOL\_E_{subject,j} + \beta_5 sexF{:}groupPOL\_L_{subject,j} + \mathbf{b_1} cohort + \mathbf{b_2} litter\ size + \epsilon_{subject,j}$

Y= outcome measures (i.e. blurred absolute Jacobian determinants); $\beta_i$= fixed effect coefficient; $\beta_0$ = equation intercept; **b** = random predictor; **ϵ** = random error; **j** = repeated measure per subject; **:** = interaction; POL_E = early polyI:C group relative to

SAL as reference; POL_L = late polyI:C group relative to SAL as reference; SexF = female sex relative to male as reference

As an exploratory analysis, we investigated whether there were any volume differences in the organs due to GD9 or 17 MIA exposure. We ran the same model as above on the voxels within the organ cavity defined manually with the same models described above.

#### 2. Supplementary results

##### 2.1 Poly I:C injection does increase pro-inflammatory cytokines

We observed an increase in levels of pro-inflammatory cytokines IL-6, IL-1β**,** and IL-10, but not TNF-α, in a separate cohort of pregnant dams 3-hours post poly I:C injection with our first batch on GD 9 relative to saline control on GD 9. For the second batch of poly I:C injected on GD 9, all 4 pro-inflammatory cytokines were increased (IL-6, IL-1β, TNF-α) 3 hours post-injection, but the anti-inflammatory cytokine IL-10 was not.

Exposure to our first batch of poly I:C (batch 1) on GD 17 increased levels of pro-inflammatory cytokines IL-6 relative to all but one SAL L dam which had exceptionally high IL-6 values. No differences were observed for TNF-α, IL-1β, or IL-10 relative to GD 17 saline controls. Our second batch of poly I:C similarly only increased IL-6, and had no effect on the other cytokine levels, however, for IL-1β and IL-10 values were below the detection threshold (<0.64) (**Supplementary Table 1)**.

**Supplementary Table 1.** Maternal serum cytokine levels for our 4 treatment groups, mean [range]; * below detection range. Batch 1 of poly I:C was used for cohort 1 and batch 2 of poly I:C for cohort 2.

| | **IL-1β (pg/ml)** | **IL-6 (pg/ml)** | **IL-10 (pg/ml)** | **TNF-α (pg/ml)** |
|---|---|---|---|---|
| **SAL E (n=5)** | 21.58 [0.64-96.62] | 45.54 [30.58-56.59] | 21.86 [0.64-56.22] | 14.30 [0.64-26.05] |
| **SAL L (n=3)** | 4.62 [0.64-6.88] | 746.33 [22.2-2192.47] | 42.13 [8.30-83.37] | 17.76 [5.99-39.59] |
| **POL E batch 1 (n=3)** | 11.56 [9.80-12.95] | 3578.84 [3624.87-3740.81] | 44.05 [36.93-49.70] | 8.20 [7.91-8.35] |
| **POL L batch 1 (n=3)** | 5.45 [5.00-6.34] | 1298.69 [54.36-3173.25] | 16.73 [13.67-19.58] | 20.35 [14.21-31.18] |
| **POL E batch 2 (n=4)** | 16.26 [0.64-25.4] | 14086.22 [1000.00-18033.31] | 76.79 [35.18-174.44] | 69.05 [7.02-131.31] |
| **POL L batch 2 (n=4)** | 0.64* [0.64-0.64] | 1503.49 [0.64-1000] | 0.64 [0.64-0.64] | 4.95 [0.64-14.98] |

#### 2.2 MRI results

##### 2.2.1 Difference between POL E and POL L

Significant brain volume differences were also observed between the POL E and POL L offspring ((t=3.590, <1%FDR) wherein the POL L offspring also had larger brain volume than POL E Offspring (not just SAL) in may similar regions such as, but not limited to, the left caudate, the bilateral dorsal hippocampus, the bilateral medial preoptic area, the bilateral septal nucleus (posterior), the anterior commissure, external capsule, and centromedian thalamus (as well as other thalamic nuclei). The POL L offspring had smaller volume than POL E offspring in the anterior septal nucleus, the cingulum/hippocampus, with mixed results in the cerebellum (some increased regions and some decreased).

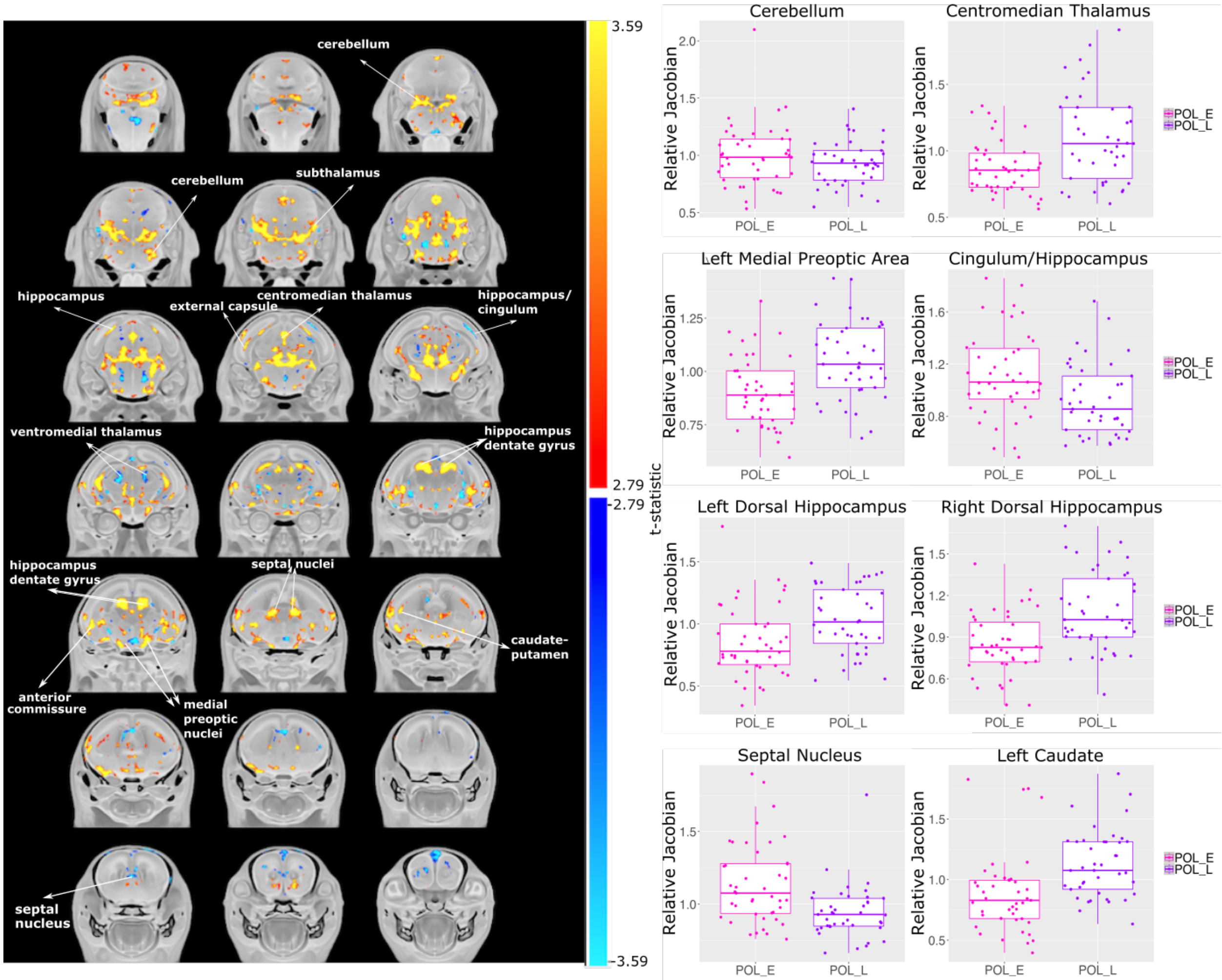


**Supplementary Figure 1.** Neuroanatomical differences due to early (GD9) or late (GD17) MIA-exposure in the GD 18 embryo mouse brain. **A.** t-statistic map of group (POL L vs POL E) thresholded at 5% (bottom, t=2.79) and 1% FDR (top, t=3.59) overlaid on the study average. **B.** Boxplots of peak voxels (voxels within a region of volume change showing largest effect) selected from regions of interest highlighted in white text in **A**. For all boxplots, the relative Jacobian determinants are plotted on the y-axis as in Figure 2 and 3.

##### 2.2.2. No volumetric differences in the embryo organs

No significant differences were observed in the volume of the organ cavity for POL E or POL L exposed embryos at GD 18 (data not shown). The average for the whole-body registration is presented below in **Supplementary figure 2.**

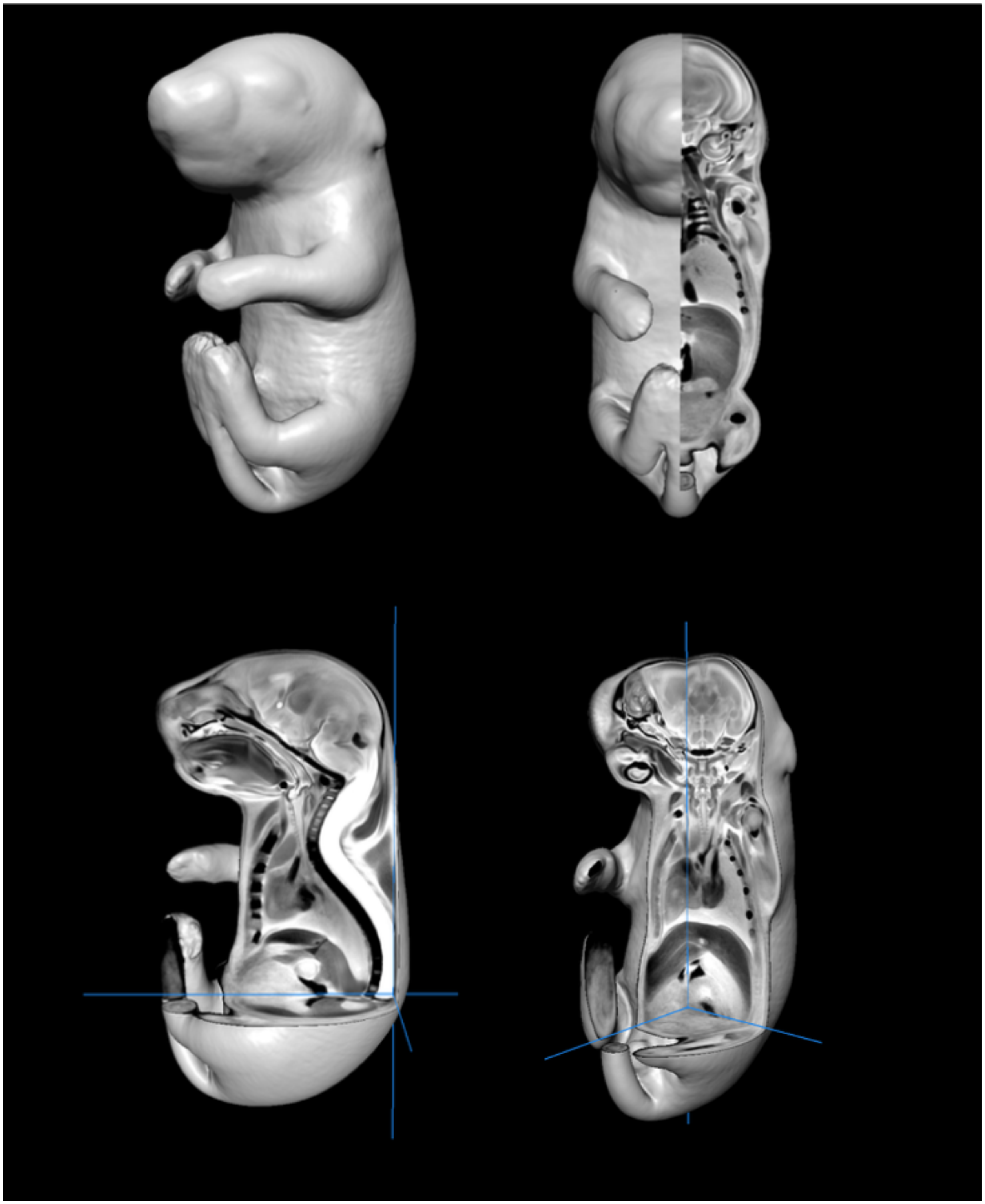

**Supplementary figure 2.** Representative 3D view of whole-body nonlinear average for the GD 18 embryo (40 μm$^3$ resolution).

#### 2.3 Electron microscopy results

##### 2.3.1 No differences in total cell density

As discussed in the main paper section 3.2, there were no differences in total cell density (chi-squared = 0.68038, df = 2, p-value = 0.7116; **Supplementary figure 3**). Further representative sections displayed in **Supplementary figure 4** per cell type per group.

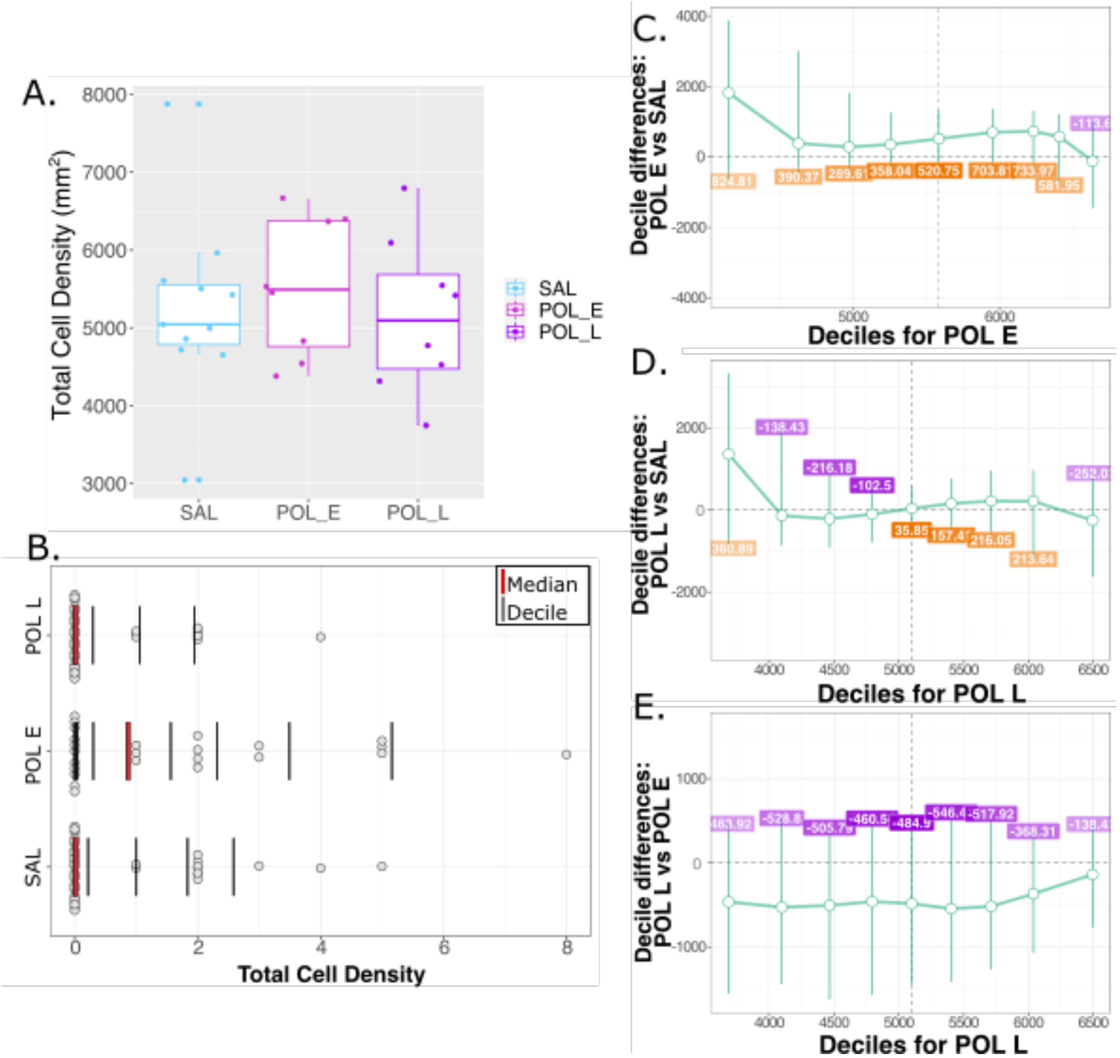


**Supplementary figure 3.** No differences in total cell density between groups. **A.** Boxplot showing group differences between SAL, POL E, and POL L offspring total cell density in the dorsal hippocampus. **B.** Distribution plot in which the red line identifies the median of the data, while each black bar denotes a decile of distribution. **C, D, and E** show results from the percentile bootstrapping technique applied to identify the difference in decile between the POL E and SAL **(C)**, POL L and SAL **(D)**, and POL E and POL L **(E)**.

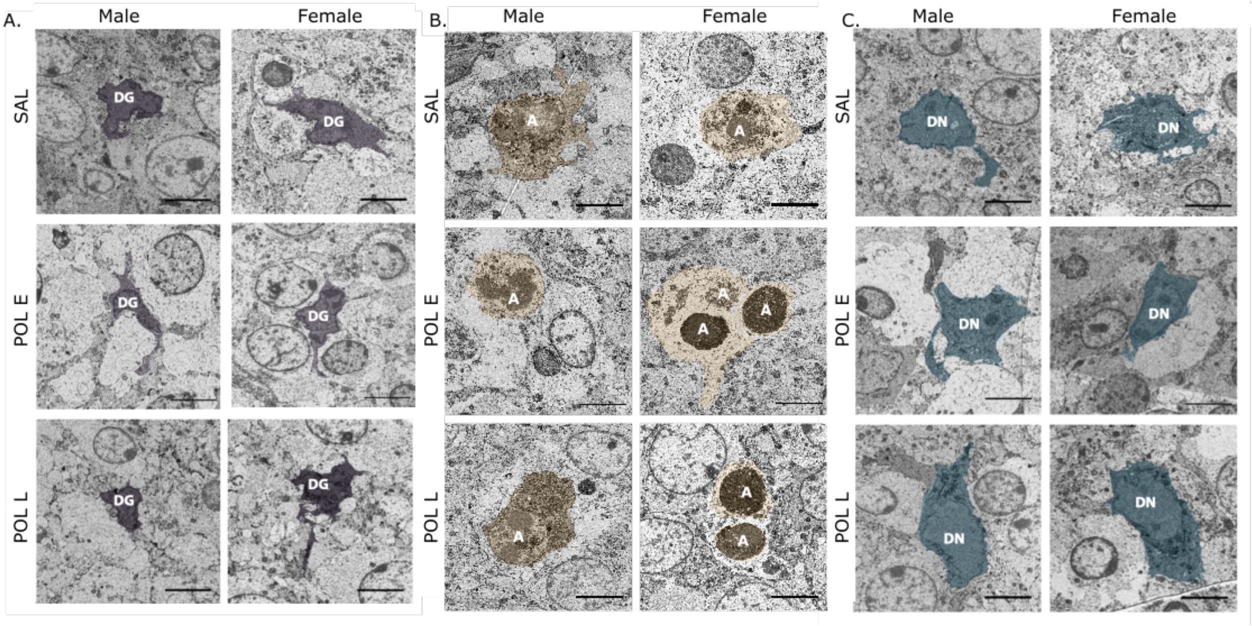


**Supplementary figure 4.** Image acquired by scanning electron microscopy (25 nm resolution) in the dorsal hippocampus from representative offspring (equivalent to coronal slice 14 from **Figure 4A**) highlighting dark glial cells (DG) in the left panel, apoptotic cells (A) in the middle panel, and dark neurons (DN) in the rightmost panel. To aid with visibility, apoptotic cells arepseudo-coloured in pale orange and dark neurons in dark blue. Scale bar equivalent to 5 μm.

##### 2.3.2 Sex differences in distribution

As was done for the MRI data, sex differences were investigated as a secondary, exploratory analysis. For total cell density there were largely no differences in distribution for POL E males (apart from lower density at the first decile for POL E males, p=0.05) nor POL L males relative to SAL (apart from lower density at the first decile for POL L males, p=0.019), nor for POL L males relative to POL E males. POL E females did have slightly higher density than both SAL (deciles 1-6, p<0.038) and POL L females (deciles 1-4, p<0.05) while POL L females were no different than SAL females (**Supplementary Tables 14-19; Supplementary Figure 5AD).**

A greater sex-dependence emerged with the dark glial cell density, wherein POL L females had significantly higher density than SAL and POL E females across all deciles of the distribution (p<0.015). Further, POL E females had even lower levels of glial cell density than SAL across higher deciles of the distribution (deciles 5-9, p<0.041). In contrast, POL L males had lower density than SAL males at lower deciles of distribution (deciles 1-6, p<0.048), and than POL E males at the first decile (p=0.29). Similar to the females, POL E males also had lower density than SAL at higher deciles (deciles 5-9, p<0.029) (**Supplementary Tables 20-26; Supplementary Figure 5EH)**.

For dark neurons, both POL E males had lower density than SAL at the higher deciles of distribution (deciles 5-9, p<0.037), while POL L males had lower density than

SAL at the lower deciles (deciles 1-6, $p<0.008$). There were no differences in distribution between POL L and POL E males, nor for any of the female pairings (**Supplementary Tables 27-32; Supplementary Figure 5IL)**.

Finally, for apoptotic cells, POL E females had significantly higher density across most deciles of the distribution relative to SAL (deciles 107, $p<0.044$), and across all deciles relative to POL L ($p<0.005$). Males instead showed no differences for any pairwise comparisons (**Supplementary Tables 33-37; Supplementary Figure 5MP).**

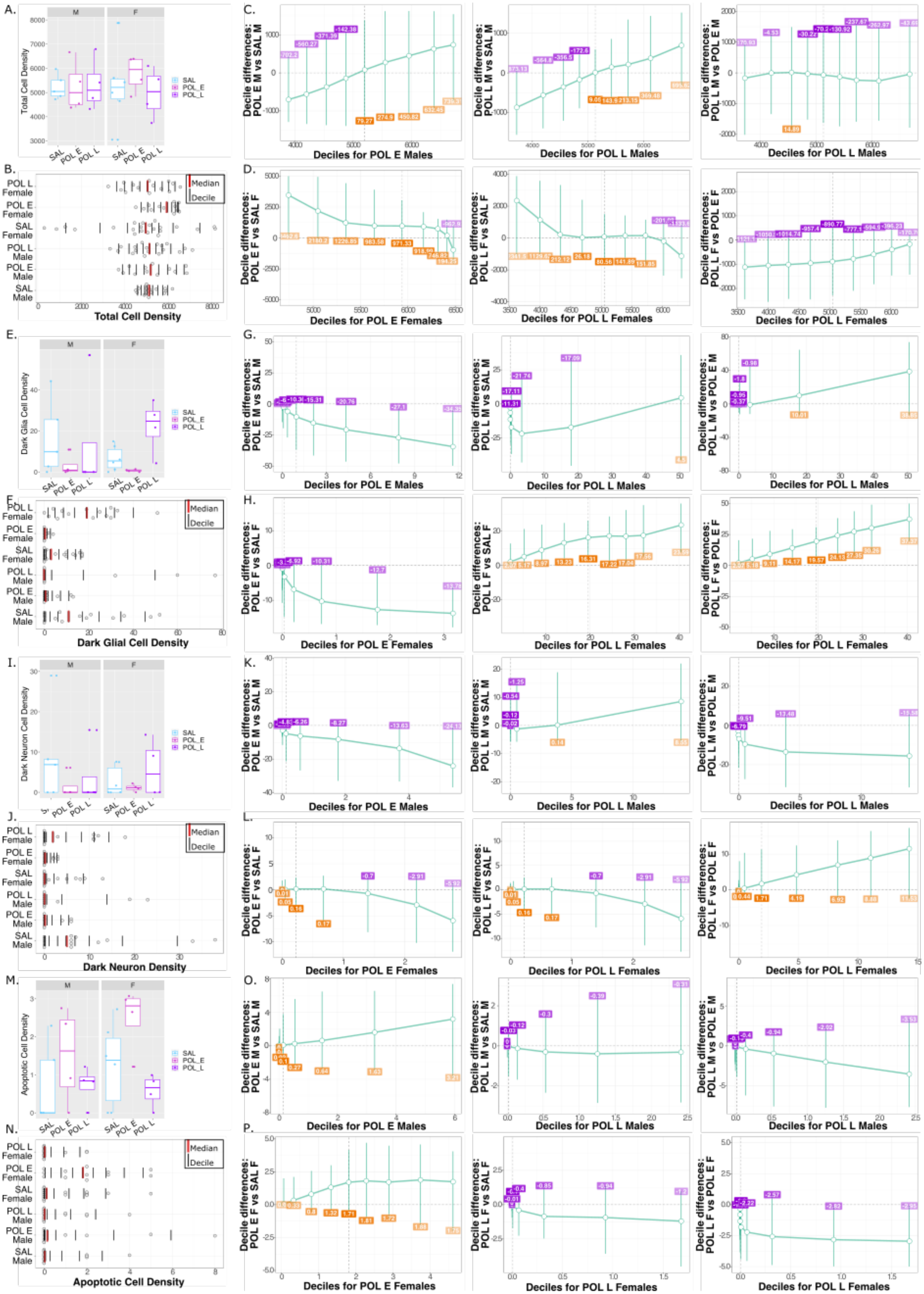
A.
Total Cell Density
SAL
POL_E
POL_L
B.
Median
Decile
Total Cell Density
C.
Decile differences: POL E M vs SAL M
Deciles for POL E Males
Decile differences: POL L M vs SAL M
Deciles for POL L Males
Decile differences: POL L M vs POL E M
Deciles for POL L Males
D.
Decile differences: POL E F vs SAL F
Deciles for POL E Females
Decile differences: POL L F vs SAL F
Deciles for POL L Females
Decile differences: POL L F vs POL E F
Deciles for POL L Females
E.
Dark Glia Cell Density
F.
Dark Glial Cell Density
G.
Deciles for POL E Males
Deciles for POL L Males
Deciles for POL L Males
H.
Deciles for POL E Females
Deciles for POL L Females
Deciles for POL L Females
I.
Dark Neuron Cell Density
J.
Dark Neuron Density
K.
Deciles for POL E Males
Deciles for POL L Males
Deciles for POL L Males
L.
Deciles for POL E Females
Deciles for POL L Females
Deciles for POL L Females
M.
Apoptotic Cell Density
N.
Apoptotic Cell Density
O.
Deciles for POL E Males
Deciles for POL L Males
Deciles for POL L Males
P.
Deciles for POL E Females
Deciles for POL L Females
Deciles for POL L Females

**Supplementary figure 5.** Differences in distribution of dark glial, dark neuron, and apoptotic cell density per sex and per group. **A.** Boxplot showing total cell density (per mouse) per sex per group (n=3-4/sex/group). **B**. Distribution of total cell density for all hippocampal slices per animal for each sex/group. The red line identifies the median of the data, while each black bar denotes a decile of distribution. A percentile bootstrapping technique applied to identify the difference in decile between the POL E and SAL, POL L and SAL, and POL E and POL L for males (**C)** and females (**D)** showing generally no differences. **E.** Boxplot showing dark glial cell density (per mouse) per sex per group (n=3-4/sex/group). **F**. Distribution of dark glial cell density for all hippocampal slices per animal for each sex/group. Difference in decile between all groups for males (**G**) and females (**H**) wherein POL L females had higher density than SAL and POL E, and POL E had lower density than SAL and POL L. **I.** Boxplot showing dark neuron density (per mouse) per sex per group (n=3-4/sex/group). **J.** Distribution of dark neuron density for all hippocampal slices per animal for each sex/group.. Difference in decile between all groups for males (**K**) and females (**L**). **M.** Boxplot showing apoptotic cell density (per mouse) per sex per group (n=3-4/sex/group). **N.** Distribution of apoptotic cell density for all hippocampal slices per animal for each sex/group. Difference in decile between all groups for males (**O**) and females (**P**).

#### Supplementary tables

For supplementary tables **2-37**, CI (confidence interval), p crit (uncorrected p-value), p-value (bootstrap corrected p-value); SAL (saline); POL E (early poly I:C group); POL L (late poly I:C group).

##### Supplementary Table 2. Total Cell Density POL E vs SAL difference

| Decile | POL E | SAL | Difference | CI lower | CI upper | p crit | p-value |
|---|---|---|---|---|---|---|---|
| 1 | 4151.408 | 2326.602 | 1824.8056 | -685.2851 | 3881.0665 | 0.00277778 | 0.092 |
| 2 | 4627.183 | 4236.812 | 390.3706 | -557.2165 | 3007.9732 | 0.00833333 | 0.358 |
| 3 | 4972.235 | 4682.629 | 289.6055 | -464.8035 | 1833.0812 | 0.00555556 | 0.286 |
| 4 | 5255.654 | 4897.611 | 358.0427 | -386.0938 | 1266.3072 | 0.00416667 | 0.190 |
| 5 | 5584.768 | 5064.015 | 520.7527 | -381.3019 | 1342.328 | 0.00238095 | 0.083 |
| 6 | 5950.95 | 5247.137 | 703.8125 | -377.1873 | 1373.1058 | 0.00208333 | 0.066 |

| | | | | | | | |
|---|---|---|---|---|---|---|---|
| 7 | 6230.605 | 5496.634 | 733.9711 | -382.9449 | 1320.4004 | 0.00185185 | 0.045 |
| 8 | 6407.253 | 5825.298 | 581.955 | -793.8431 | 1215.9668 | 0.00333333 | 0.136 |
| 9 | 6633.453 | 6747.055 | -113.602 | -1442.54 | 964.3226 | 0.01666667 | 0.892 |

#### Supplementary Table 3. Total Cell Density POL L vs SAL difference

| Decile | POL L | SAL | Difference | CI lower | CI upper | p crit | p-value |
|---|---|---|---|---|---|---|---|
| 1 | 3687.489 | 2326.602 | 1360.88663 | -933.4781 | 3332.1491 | 0.00555556 | 0.278 |
| 2 | 4098.382 | 4236.812 | -138.4302 | -861.9588 | 2023.1686 | 0.025 | 0.848 |
| 3 | 4466.444 | 4682.629 | -216.18463 | -917.2317 | 1036.4123 | 0.00714286 | 0.537 |
| 4 | 4795.114 | 4897.611 | -102.49719 | -788.0476 | 604.2014 | 0.0125 | 0.764 |
| 5 | 5099.865 | 5064.015 | 35.85004 | -488.4449 | 579.8772 | 0.05 | 0.894 |
| 6 | 5404.544 | 5247.137 | 157.40696 | -617.1548 | 763.2366 | 0.01 | 0.648 |
| 7 | 5712.688 | 5496.634 | 216.05434 | -729.4657 | 961.1043 | 0.00625 | 0.495 |
| 8 | 6038.94 | 5825.298 | 213.64173 | -1196.4319 | 968.0088 | 0.00833333 | 0.623 |
| 9 | 6495.022 | 6747.055 | -252.03283 | -1624.8705 | 897.5734 | 0.01666667 | 0.732 |

#### Supplementary Table 4. Total Cell Density POL L vs POL E difference

| Decile | POL L | POL E | Difference | CI lower | CI upper | p crit | p-value |
|---|---|---|---|---|---|---|---|
| 1 | 3687.489 | 4151.408 | -463.9189 | -1557.1103 | 465.3654 | 0.00185185 | 0.112 |
| 2 | 4098.382 | 4627.183 | -528.8008 | -1442.7045 | 531.7885 | 0.00416667 | 0.155 |
| 3 | 4466.444 | 4972.235 | -505.7901 | -1625.6035 | 419.6274 | 0.00277778 | 0.151 |

| | | | | | | | |
|---|---|---|---|---|---|---|---|
| **4** | 4795.114 | 5255.654 | -460.5399 | -1571.5615 | 525.1342 | 0.00333333 | 0.160 |
| **5** | 5099.865 | 5584.768 | -484.9026 | -1446.6147 | 476.4443 | 0.00555556 | 0.195 |
| **6** | 5404.544 | 5950.95 | -546.4056 | -1416.1051 | 600.4311 | 0.00238095 | 0.189 |
| **7** | 5712.688 | 6230.605 | -517.9168 | -1268.8219 | 582.2052 | 0.00208333 | 0.136 |
| **8** | 6038.94 | 6407.253 | -368.3133 | -1071.0121 | 375.3764 | 0.00833333 | 0.184 |
| **9** | 6495.022 | 6633.453 | -138.4309 | -772.0971 | 459.7784 | 0.01666667 | 0.625 |

#### Supplementary Table 5. Total Dark Glial Cell Density POL E vs SAL difference

| Decile | POL E | SAL | Difference | CI lower | CI upper | p crit | p-value |
|---|---|---|---|---|---|---|---|
| **1** | 1.56E-08 | 1.65E-05 | -1.65E-05 | -0.117288 | 0.0008025 | 0.00625 | 0.283 |
| **2** | 7.42E-06 | 6.43E-03 | -6.43E-03 | -2.801359 | 0.07893029 | 0.003125 | 0.201 |
| **3** | 6.21E-04 | 2.32E-01 | -2.32E-01 | -6.841321 | 0.29481655 | 0.00208333 | 0.109 |
| **4** | 1.56E-02 | 1.88E+00 | -1.87E+00 | -10.613885 | 0.4750728 | 0.0015625 | 0.067 |
| **5** | 1.48E-01 | 5.73E+00 | -5.58E+00 | -14.32128 | 0.93706151 | 0.00125 | 0.020 |
| **6** | 6.36E-01 | 1.01E+01 | -9.51E+00 | -17.339485 | -0.2218018 | 0.00104167 | 0.001 |
| **7** | 1.59E+00 | 1.44E+01 | -1.28E+01 | -23.66778 | -2.6836544 | 0.00078125 | <0.001 |
| **8** | 3.54E+00 | 1.85E+01 | -1.50E+01 | -39.659531 | -2.5981535 | 0.00089286 | <0.001 |
| **9** | 8.55E+00 | 3.11E+01 | -2.25E+01 | -45.872087 | -4.508818 | 0.00069444 | <0.001 |

#### Supplementary Table 6. Total Dark Glial Cell Density POL L vs SAL difference

| Decile | POL L | SAL | Difference | CI lower | CI upper | p crit | p-value |
|---|---|---|---|---|---|---|---|
| 1 | 7.35E-07 | 1.56E-08 | 7.20E-07 | -0.0091239 | 0.07514775 | 6.94E-04 | 0.562 |
| 2 | 4.50E-04 | 7.42E-06 | 4.42E-04 | -0.3304835 | 1.09187281 | 3.47E-04 | 0.478 |
| 3 | 3.15E-02 | 6.21E-04 | 3.08E-02 | -0.3715421 | 8.08383966 | 2.31E-04 | 0.339 |
| 4 | 5.27E-01 | 1.56E-02 | 5.11E-01 | -1.2459148 | 13.9375972 | 1.74E-04 | 0.241 |
| 5 | 3.24E+00 | 1.48E-01 | 3.09E+00 | -2.386622 | 24.4702609 | 1.39E-04 | 0.124 |
| 6 | 1.03E+01 | 6.36E-01 | 9.67E+00 | -2.377137 | 28.7006809 | 1.16E-04 | 0.043 |
| 7 | 2.06E+01 | 1.59E+00 | 1.90E+01 | -1.7454353 | 39.9903739 | 9.92E-05 | 0.008 |
| 8 | 2.97E+01 | 3.54E+00 | 2.62E+01 | 1.16311037 | 56.2422815 | 8.68E-05 | <0.001 |
| 9 | 4.54E+01 | 8.55E+00 | 3.68E+01 | 7.18141072 | 70.1908463 | 7.72E-05 | <0.001 |

#### Supplementary Table 7. Total Dark Glial Cell Density POL L vs POL E difference

| Decile | POL L | POL E | Difference | CI lower | CI upper | p crit | p-value |
|---|---|---|---|---|---|---|---|
| 1 | 7.35E-07 | 1.65E-05 | -1.58E-05 | -0.0408415 | 0.01028092 | 0.01666667 | 0.634 |
| 2 | 4.50E-04 | 6.43E-03 | -5.98E-03 | -1.2774581 | 0.50064786 | 0.01 | 0.597 |
| 3 | 3.15E-02 | 2.32E-01 | -2.01E-01 | -5.6629882 | 4.09638211 | 0.00833333 | 0.531 |
| 4 | 5.27E-01 | 1.88E+00 | -1.36E+00 | -9.0806618 | 8.04585178 | 0.0125 | 0.542 |
| 5 | 3.24E+00 | 5.73E+00 | -2.49E+00 | -11.367416 | 12.7351975 | 0.025 | 0.683 |
| 6 | 1.03E+01 | 1.01E+01 | 1.55E-01 | -11.178672 | 15.8678074 | 0.05 | 0.968 |
| 7 | 2.06E+01 | 1.44E+01 | 6.23E+00 | -13.049187 | 24.0356019 | 0.00714286 | 0.435 |

| | | | | | | | |
|---|---|---|---|---|---|---|---|
| **8** | 2.97E+01 | 1.85E+01 | 1.12E+01 | -14.425163 | 34.1453 | 0.00555556 | 0.181 |
| **9** | 4.54E+01 | 3.11E+01 | 1.43E+01 | -15.408127 | 48.9480906 | 0.00625 | 0.255 |

#### Supplementary Table 8. Total Dark Neuron Density POL E vs SAL difference

| Decile | POL E | SAL | Difference | CI lower | CI upper | p crit | p-value |
|---|---|---|---|---|---|---|---|
| **1** | 9.72E-11 | 4.80E-10 | -3.83E-10 | -4.15E-05 | 1.11E-05 | 0.025 | 0.851 |
| **2** | 1.15E-07 | 2.01E-06 | -1.90E-06 | -1.95E-02 | 5.61E-03 | 0.0125 | 0.657 |
| **3** | 2.37E-05 | 6.84E-04 | -6.60E-04 | -7.39E-01 | 1.49E-01 | 0.00833333 | 0.554 |
| **4** | 1.43E-03 | 4.08E-02 | -3.94E-02 | -3.36E+00 | 5.12E-01 | 0.00625 | 0.438 |
| **5** | 3.13E-02 | 6.07E-01 | -5.76E-01 | -5.59E+00 | 1.26E+00 | 0.005 | 0.299 |
| **6** | 2.81E-01 | 2.83E+00 | -2.55E+00 | -6.92E+00 | 1.87E+00 | 0.00416667 | 0.191 |
| **7** | 1.15E+00 | 5.58E+00 | -4.44E+00 | -9.81E+00 | 1.42E+00 | 0.00357143 | 0.051 |
| **8** | 2.59E+00 | 8.05E+00 | -5.47E+00 | -2.00E+01 | 4.47E-01 | 0.003125 | 0.010 |
| **9** | 4.56E+00 | 1.55E+01 | -1.09E+01 | -3.14E+01 | -1.82E+00 | 0.00277778 | <0.001 |

#### Supplementary Table 9. Total Dark Neuron Density POL L vs SAL difference

| Decile | POL L | SAL | Difference | CI lower | CI upper | p crit | p-value |
|---|---|---|---|---|---|---|---|
| **1** | 6.46E-14 | 4.80E-10 | -4.80E-10 | -1.61E-04 | 1.36E-07 | 0.0125 | 0.3585 |
| **2** | 7.49E-10 | 2.01E-06 | -2.01E-06 | -2.45E-02 | 1.10E-04 | 0.01 | 0.330 |
| **3** | 9.04E-07 | 6.84E-04 | -6.83E-04 | -6.13E-01 | 2.90E-02 | 0.00714286 | 0.339 |
| **4** | 2.21E-04 | 4.08E-02 | -4.06E-02 | -3.24E+00 | 4.67E-01 | 0.00625 | 0.275 |

| | | | | | | | |
|---|---|---|---|---|---|---|---|
| **5** | 1.46E-02 | 6.07E-01 | -5.93E-01 | -5.25E+00 | 2.81E+00 | 0.00555556 | 0.287 |
| **6** | 3.15E-01 | 2.83E+00 | -2.52E+00 | -6.90E+00 | 5.72E+00 | 0.00833333 | 0.304 |
| **7** | 2.51E+00 | 5.58E+00 | -3.08E+00 | -8.49E+00 | 6.47E+00 | 0.01666667 | 0.479 |
| **8** | 8.10E+00 | 8.05E+00 | 4.66E-02 | -9.50E+00 | 7.19E+00 | 0.05 | 0.936 |
| **9** | 1.50E+01 | 1.55E+01 | -4.46E-01 | -1.90E+01 | 9.97E+00 | 0.025 | 0.860 |

#### Supplementary Table 10. Total Dark Neuron Density POL L vs POL E difference

| **Decile** | **POL L** | **POL E** | **Difference** | **CI lower** | **CI upper** | **p crit** | **p-value** |
|---|---|---|---|---|---|---|---|
| **1** | 6.46E-14 | 9.72E-11 | -9.71E-11 | -0.0011336 | 4.43E-06 | 0.00039683 | 0.533 |
| **2** | 7.49E-10 | 1.15E-07 | -1.14E-07 | -0.017834 | 7.33E-04 | 0.00055556 | 0.588 |
| **3** | 9.04E-07 | 2.37E-05 | -2.28E-05 | -0.2128114 | 6.68E-02 | 0.00069444 | 0.706 |
| **4** | 2.21E-04 | 1.43E-03 | -1.21E-03 | -1.1878761 | 9.77E-01 | 0.00092593 | 0.785 |
| **5** | 1.46E-02 | 3.13E-02 | -1.67E-02 | -2.1454795 | 3.32E+00 | 0.00277778 | 0.910 |
| **6** | 3.15E-01 | 2.81E-01 | 3.39E-02 | -2.7517723 | 8.25E+00 | 0.00138889 | 0.905 |
| **7** | 2.51E+00 | 1.15E+00 | 1.36E+00 | -3.808122 | 1.18E+01 | 0.00046296 | 0.576 |
| **8** | 8.10E+00 | 2.59E+00 | 5.51E+00 | -5.1950725 | 1.67E+01 | 0.00034722 | 0.230 |
| **9** | 1.50E+01 | 4.56E+00 | 1.05E+01 | -4.0620577 | 1.91E+01 | 0.00030864 | 0.016 |

#### Supplementary Table 11. Total Apoptotic Cell Density POL E vs SAL difference

| Decile | POL E | SAL | Difference | CI lower | CI upper | p crit | p-value |
|---|---|---|---|---|---|---|---|
| 1 | 2.98E-05 | 3.20E-14 | 2.98E-05 | -4.43E-07 | 0.07461499 | 0.00069444 | 0.013 |
| 2 | 2.84E-03 | 6.63E-10 | 2.84E-03 | -1.04E-03 | 0.85163803 | 0.00078125 | 0.020 |
| 3 | 5.05E-02 | 1.04E-06 | 5.05E-02 | -3.11E-02 | 1.35800997 | 0.00089286 | 0.028 |
| 4 | 3.03E-01 | 2.62E-04 | 3.03E-01 | -1.75E-01 | 2.33253458 | 0.00104167 | 0.039 |
| 5 | 8.67E-01 | 1.45E-02 | 8.53E-01 | -7.87E-01 | 2.84572975 | 0.0015625 | 0.064 |
| 6 | 1.56E+00 | 2.16E-01 | 1.35E+00 | -1.01E+00 | 3.73112757 | 0.003125 | 0.124 |
| 7 | 2.31E+00 | 1.00E+00 | 1.31E+00 | -1.04E+00 | 4.21302706 | 0.00625 | 0.121 |
| 8 | 3.49E+00 | 1.84E+00 | 1.66E+00 | -9.74E-01 | 4.89217444 | 0.00208333 | 0.083 |
| 9 | 5.16E+00 | 2.58E+00 | 2.58E+00 | -1.13E+00 | 5.81488105 | 0.00125 | 0.057 |

#### Supplementary Table 12. Total Apoptotic Cell Density POL L vs SAL difference

| Decile | POL L | SAL | Difference | CI lower | CI upper | p crit | p-value |
|---|---|---|---|---|---|---|---|
| 1 | 6.66E-16 | 3.20E-14 | -3.13E-14 | -1.62E-08 | 7.86E-10 | 0.05 | 0.727 |
| 2 | 1.39E-11 | 6.63E-10 | -6.49E-10 | -3.36E-05 | 7.97E-06 | 0.025 | 0.654 |
| 3 | 2.78E-08 | 1.04E-06 | -1.01E-06 | -9.36E-03 | 1.07E-03 | 0.01666667 | 0.654 |
| 4 | 1.10E-05 | 2.62E-04 | -2.51E-04 | -2.28E-01 | 3.02E-02 | 0.0125 | 0.626 |
| 5 | 1.11E-03 | 1.45E-02 | -1.34E-02 | -8.71E-01 | 2.40E-01 | 0.01 | 0.534 |
| 6 | 3.25E-02 | 2.16E-01 | -1.83E-01 | -1.64E+00 | 7.34E-01 | 0.00833333 | 0.470 |
| 7 | 2.99E-01 | 1.00E+00 | -7.05E-01 | -1.93E+00 | 1.17E+00 | 0.00714286 | 0.398 |

| | | | | | | | |
|---|---|---|---|---|---|---|---|
| **8** | 1.06E+00 | 1.84E+00 | -7.78E-01 | -2.32E+00 | 1.25E+00 | 0.00555556 | 0.288 |
| **9** | 1.95E+00 | 2.58E+00 | -6.31E-01 | -3.03E+00 | 1.40E+00 | 0.00625 | 0.353 |

#### Supplementary Table 13. Total Apoptotic Cell Density POL L vs POL E difference

| Decile | POL L | POL E | Difference | CI lower | CI upper | p crit | p-value |
|---|---|---|---|---|---|---|---|
| **1** | 6.66E-16 | 2.98E-05 | -2.98E-05 | -0.3070457 | 3.17E-08 | 0.00013889 | 0.006 |
| **2** | 1.39E-11 | 2.84E-03 | -2.84E-03 | -0.9555023 | 6.64E-06 | 0.00015625 | 0.003 |
| **3** | 2.78E-08 | 5.05E-02 | -5.05E-02 | -1.8488396 | 9.25E-04 | 0.0003125 | 0.008 |
| **4** | 1.10E-05 | 3.03E-01 | -3.03E-01 | -2.4652326 | 2.24E-02 | 0.00025 | 0.009 |
| **5** | 1.11E-03 | 8.67E-01 | -8.66E-01 | -2.8238152 | 1.43E-01 | 0.000625 | 0.015 |
| **6** | 3.25E-02 | 1.56E+00 | -1.53E+00 | -3.9222594 | 4.22E-01 | 0.00125 | 0.020 |
| **7** | 2.99E-01 | 2.31E+00 | -2.01E+00 | -4.9137293 | 7.29E-01 | 0.00020833 | 0.011 |
| **8** | 1.06E+00 | 3.49E+00 | -2.43E+00 | -6.3036623 | 2.18E-01 | 0.00041667 | 0.007 |
| **9** | 1.95E+00 | 5.16E+00 | -3.21E+00 | -6.6126179 | 1.50E+00 | 0.00017857 | 0.014 |

#### Supplementary Table 14. Total Cell Density POL E vs SAL Males difference

| Decile | POL E Males | SAL Males | Difference | CI lower | CI upper | p crit | p-value |
|---|---|---|---|---|---|---|---|
| **1** | 3890.556 | 4592.76 | -702.20394 | -1305.4195 | 563.4912 | 9.92E-05 | 0.050 |
| **2** | 4198.377 | 4758.65 | -560.27311 | -1263.0703 | 1046.9528 | 1.28E-04 | 0.165 |

| | | | | | | | |
|---|---|---|---|---|---|---|---|
| **3** | 4541.701 | 4913.09 | -371.38933 | -1431.1522 | 1022.9294 | 2.23E-04 | 0.402 |
| **4** | 4875.572 | 5017.954 | -142.38194 | -1134.8485 | 1126.6137 | 4.46E-04 | 0.723 |
| **5** | 5198.289 | 5119.018 | 79.27151 | -1152.8401 | 1482.5871 | 8.93E-04 | 0.878 |
| **6** | 5551.005 | 5276.107 | 274.89765 | -1102.0542 | 1564.0392 | 2.98E-04 | 0.571 |
| **7** | 5958.77 | 5507.953 | 450.81698 | -889.3568 | 1638.6579 | 1.79E-04 | 0.400 |
| **8** | 6384.625 | 5752.175 | 632.4499 | -915.8302 | 1525.3192 | 1.49E-04 | 0.204 |
| **9** | 6719.294 | 5979.987 | 739.30689 | -917.1017 | 1659.7137 | 1.12E-04 | 0.082 |

#### Supplementary Table 15. Total Cell Density POL E vs SAL Females difference

| Decile | POL E Females | SAL Females | Difference | CI lower | CI upper | p crit | p-value |
|---|---|---|---|---|---|---|---|
| **1** | 4740.61 | 1278.007 | 3462.6034 | 237.65558 | 5054.735 | 1.24E-05 | <0.001 |
| **2** | 5050.65 | 2870.448 | 2180.2024 | -16.12843 | 4808.611 | 1.40E-05 | 0.002 |
| **3** | 5343.483 | 4116.629 | 1226.8536 | -208.66979 | 4457.246 | 1.59E-05 | 0.004 |
| **4** | 5647.352 | 4663.776 | 983.576 | -246.80128 | 4170.709 | 1.86E-05 | 0.013 |
| **5** | 5938.522 | 4967.196 | 971.3257 | -348.42508 | 2812.317 | 2.23E-05 | 0.018 |
| **6** | 6165.649 | 5246.656 | 918.9926 | -870.52609 | 2246.009 | 2.79E-05 | 0.038 |
| **7** | 6315.302 | 5568.478 | 746.8239 | -1895.1188 | 1546.637 | 3.72E-05 | 0.186 |
| **8** | 6413.841 | 6219.588 | 194.2528 | -2279.906 | 1330.446 | 1.12E-04 | 0.803 |
| **9** | 6478.831 | 7441.747 | -962.9169 | -1905.2061 | 1109.814 | 5.58E-05 | 0.296 |

#### Supplementary Table 16. Total Cell Density POL L vs SAL Males difference

| Decile | POL L Males | SAL Males | Difference | CI lower | CI upper | p crit | p-value |
|---|---|---|---|---|---|---|---|
| 1 | 3719.626 | 4592.76 | -873.13456 | -1565.047 | 104.1645 | 0.00555556 | 0.019 |
| 2 | 4193.846 | 4758.65 | -564.80345 | -1404.595 | 323.8518 | 0.00625 | 0.091 |
| 3 | 4556.589 | 4913.09 | -356.50166 | -1210.0169 | 397.5971 | 0.00833333 | 0.252 |
| 4 | 4845.353 | 5017.954 | -172.60043 | -894.2499 | 566.5292 | 0.01666667 | 0.555 |
| 5 | 5128.069 | 5119.018 | 9.050946 | -672.8681 | 596.2203 | 0.05 | 0.966 |
| 6 | 5420.083 | 5276.107 | 143.975209 | -700.5618 | 859.4345 | 0.025 | 0.725 |
| 7 | 5721.102 | 5507.953 | 213.149652 | -689.2292 | 1347.9808 | 0.0125 | 0.538 |
| 8 | 6121.657 | 5752.175 | 369.481884 | -575.4137 | 1439.9775 | 0.01 | 0.326 |
| 9 | 6675.606 | 5979.987 | 695.618625 | -309.4015 | 1524.7984 | 0.00714286 | 0.104 |

#### Supplementary Table 17. Total Cell Density POL L vs SAL Females difference

| Decile | POL L Females | SAL Females | Difference | CI lower | CI upper | p crit | p-value |
|---|---|---|---|---|---|---|---|
| 1 | 3619.508 | 1278.007 | 2341.50133 | -1145.09 | 3878.7923 | 0.00079365 | 0.058 |
| 2 | 4000.07 | 2870.448 | 1129.6216 | -1132.916 | 3584.8689 | 0.00102041 | 0.306 |
| 3 | 4328.748 | 4116.629 | 212.11839 | -1338.191 | 3318.5848 | 0.00119048 | 0.641 |
| 4 | 4689.957 | 4663.776 | 26.18074 | -1150.641 | 2392.5997 | 0.00357143 | 0.850 |
| 5 | 5047.756 | 4967.196 | 80.5597 | -1485.716 | 1504.3793 | 0.00238095 | 0.858 |
| 6 | 5388.548 | 5246.656 | 141.8913 | -1471.263 | 1410.6258 | 0.00178571 | 0.817 |
| 7 | 5720.328 | 5568.478 | 151.84953 | -1620.906 | 1146.9019 | 0.00714286 | 0.863 |

| | | | | | | | |
|---|---|---|---|---|---|---|---|
| 8 | 6017.608 | 6219.588 | -201.98024 | -2357.425 | 1111.491 | 0.00142857 | 0.761 |
| 9 | 6308.068 | 7441.747 | -1133.6796 | -2521.643 | 915.5942 | 0.00089286 | 0.189 |

#### Supplementary Table 18. Total Cell Density POL L vs POL E Males difference

| Decile | POL L Males | POL E Males | Difference | CI lower | CI upper | p crit | p-value |
|---|---|---|---|---|---|---|---|
| 1 | 3719.626 | 3890.556 | -170.93062 | -2012.992 | 988.9933 | 6.20E-06 | 0.577 |
| 2 | 4193.846 | 4198.377 | -4.530336 | -1340.008 | 1315.8678 | 1.40E-05 | 0.918 |
| 3 | 4556.589 | 4541.701 | 14.887671 | -1829.684 | 1491.186 | 2.79E-05 | 0.987 |
| 4 | 4845.353 | 4875.572 | -30.218496 | -1521.274 | 1266.2713 | 1.86E-05 | 0.974 |
| 5 | 5128.069 | 5198.289 | -70.220568 | -1627.329 | 1431.4893 | 1.12E-05 | 0.886 |
| 6 | 5420.083 | 5551.005 | -130.92244 | -1739.814 | 1423.9628 | 9.30E-06 | 0.804 |
| 7 | 5721.102 | 5958.77 | -237.66733 | -1587.876 | 1667.94 | 7.97E-06 | 0.707 |
| 8 | 6121.657 | 6384.625 | -262.96802 | -1554.451 | 1560.6667 | 6.98E-06 | 0.673 |
| 9 | 6675.606 | 6719.294 | -43.688261 | -1783.333 | 1625.4563 | 5.58E-05 | 0.950 |

#### Supplementary Table 19. Total Cell Density POL L vs POL E Females difference

| Decile | POL L Females | POL E Females | Difference | CI lower | CI upper | p crit | p-value |
|---|---|---|---|---|---|---|---|
| 1 | 3619.508 | 4740.61 | -1121.1021 | -2448.207 | 44.29931 | 6.20E-06 | 0.005 |
| 2 | 4000.07 | 5050.65 | -1050.5808 | -2553.967 | 229.04088 | 6.98E-06 | 0.014 |
| 3 | 4328.748 | 5343.483 | -1014.7353 | -2429.921 | 252.09046 | 7.97E-06 | 0.023 |
| 4 | 4689.957 | 5647.352 | -957.3952 | -2450.655 | 472.80904 | 9.30E-06 | 0.050 |

| | | | | | | | |
|---|---|---|---|---|---|---|---|
| 5 | 5047.756 | 5938.522 | -890.766 | -2284.841 | 657.12797 | 1.40E-05 | 0.068 |
| 6 | 5388.548 | 6165.649 | -777.1013 | -2096.116 | 446.71426 | 1.86E-05 | 0.074 |
| 7 | 5720.328 | 6315.302 | -594.9744 | -1825.747 | 528.25807 | 1.12E-05 | 0.099 |
| 8 | 6017.608 | 6413.841 | -396.2331 | -1466.23 | 579.40294 | 2.79E-05 | 0.091 |
| 9 | 6308.068 | 6478.831 | -170.7627 | -1428.37 | 316.42526 | 5.58E-05 | 0.257 |

#### Supplementary Table 20. Dark Glial Cell Density POL E vs SAL Males difference

| Decile | POL E Males | SAL Males | Difference | CI lower | CI upper | p crit | p-value |
|---|---|---|---|---|---|---|---|
| 1 | 8.38E-04 | 0.05848903 | -0.0576506 | -5.546827 | 0.2453002 | 0.00555556 | 0.325 |
| 2 | 1.45E-02 | 0.66519951 | -0.6507007 | -14.568047 | 0.9111334 | 0.00277778 | 0.169 |
| 3 | 9.78E-02 | 2.71532356 | -2.6175609 | -19.498768 | 1.6843484 | 0.00185185 | 0.113 |
| 4 | 3.70E-01 | 6.40184165 | -6.0318349 | -26.918845 | 3.1898038 | 0.00138889 | 0.068 |
| 5 | 9.70E-01 | 11.330745 | -10.360625 | -36.812142 | 3.8026146 | 0.00111111 | 0.029 |
| 6 | 2.13E+00 | 17.4356958 | -15.305585 | -41.49908 | 3.4525547 | 0.00092593 | 0.019 |
| 7 | 4.36E+00 | 25.1249898 | -20.764221 | -46.360696 | 1.9584087 | 0.00079365 | 0.007 |
| 8 | 7.93E+00 | 35.0327005 | -27.100784 | -47.848487 | -2.3724743 | 0.00069444 | <0.001 |
| 9 | 1.16E+01 | 45.9910106 | -34.348116 | -49.747092 | -3.8059681 | 0.00061728 | 0.001 |

#### Supplementary Table 21. Dark Glial Cell Density POL E vs SAL Females difference

| Decile | POL E Females | SAL Females | Difference | CI lower | CI upper | p crit | p-value |
|---|---|---|---|---|---|---|---|

| | | | | | | | |
|---|---|---|---|---|---|---|---|
| **1** | 2.67E-07 | 0.00012305 | -1.23E-04 | -0.3919406 | 0.0215986 | 6.17E-04 | 0.274 |
| **2** | 1.77E-05 | 0.00886431 | -8.85E-03 | -6.6256465 | 0.2175098 | 3.09E-04 | 0.248 |
| **3** | 4.38E-04 | 0.14457666 | -1.44E-01 | -7.2009708 | 0.4655823 | 2.06E-04 | 0.16 |
| **4** | 5.66E-03 | 0.948631 | -9.43E-01 | -10.775766 | 0.9544673 | 1.54E-04 | 0.102 |
| **5** | 4.37E-02 | 3.30487748 | -3.26E+00 | -14.76165 | 0.9190349 | 1.23E-04 | 0.041 |
| **6** | 2.17E-01 | 7.13369719 | -6.92E+00 | -16.266664 | 1.3335848 | 1.03E-04 | 0.020 |
| **7** | 7.34E-01 | 11.0482462 | -1.03E+01 | -16.838071 | 1.2533821 | 8.82E-05 | 0.007 |
| **8** | 1.77E+00 | 14.469097 | -1.27E+01 | -17.151448 | -1.2405593 | 7.72E-05 | <0.001 |
| **9** | 3.17E+00 | 16.9509635 | -1.38E+01 | -17.795357 | -5.8959528 | 6.86E-05 | <0.001 |

#### Supplementary Table 22. Dark Glial Cell Density POL L vs SAL Males difference

| **Decile** | **POL L Males** | **SAL Males** | **Difference** | **CI lower** | **CI upper** | **p crit** | **p-value** |
|---|---|---|---|---|---|---|---|
| **1** | 1.96E-11 | 0.05848903 | -0.058489 | -5.963521 | -3.61E-06 | 0.00625 | 0.001 |
| **2** | 1.83E-08 | 0.66519951 | -0.6651995 | -10.885144 | -3.00E-04 | 0.00555556 | 0.003 |
| **3** | 4.34E-06 | 2.71532356 | -2.7153192 | -17.675691 | -7.66E-03 | 0.00714286 | 0.004 |
| **4** | 3.96E-04 | 6.40184165 | -6.4014458 | -22.330719 | -2.75E-02 | 0.00833333 | 0.007 |
| **5** | 1.63E-02 | 11.330745 | -11.314395 | -29.669673 | 3.20E-01 | 0.01 | 0.014 |
| **6** | 3.30E-01 | 17.4356958 | -17.105331 | -36.697382 | 9.43E+00 | 0.0125 | 0.048 |
| **7** | 3.39E+00 | 25.1249898 | -21.739506 | -42.869066 | 2.07E+01 | 0.01666667 | 0.178 |
| **8** | 1.79E+01 | 35.0327005 | -17.090667 | -45.318803 | 3.37E+01 | 0.025 | 0.487 |

| | | | | | | | |
|---|---|---|---|---|---|---|---|
| 9 | 5.05E+01 | 45.9910106 | 4.50162926 | -40.953521 | 3.58E+01 | 0.05 | 0.882 |

**Supplementary Table 23. Dark Glial Cell Density POL L vs SAL Females difference**

| Decile | POL L Females | SAL Females | Difference | CI lower | CI upper | p crit | p-value |
|---|---|---|---|---|---|---|---|
| 1 | 2.07428 | 0.00012305 | 2.074157 | 0.06985707 | 12.83886 | 0.00833333 | <0.001 |
| 2 | 5.177012 | 0.00886431 | 5.168148 | 0.22991869 | 21.28388 | 0.00714286 | 0.005 |
| 3 | 9.1135 | 0.14457666 | 8.968923 | 1.26540404 | 23.81753 | 0.0125 | 0.004 |
| 4 | 14.175393 | 0.948631 | 13.226762 | 1.99193799 | 24.78776 | 0.025 | 0.012 |
| 5 | 19.610566 | 3.30487748 | 16.305688 | 3.86350082 | 26.17121 | 0.05 | 0.007 |
| 6 | 24.351703 | 7.13369719 | 17.218006 | 0.68907144 | 28.56021 | 0.01666667 | 0.015 |
| 7 | 28.086174 | 11.0482462 | 17.037928 | 1.49294562 | 32.33729 | 0.00625 | 0.005 |
| 8 | 32.027146 | 14.469097 | 17.558049 | 5.29958358 | 35.22357 | 0.01 | 0.001 |
| 9 | 40.542751 | 16.9509635 | 23.591788 | 7.70944969 | 36.11253 | 0.00555556 | <0.001 |

**Supplementary Table 24. Dark Glial Cell Density POL L vs POL E Males difference**

| Decile | POL L Males | POL E Males | Difference | CI lower | CI upper | p crit | p-value |
|---|---|---|---|---|---|---|---|
| 1 | 1.96E-11 | 8.38E-04 | -0.0008384 | -0.8403059 | 2.57E-04 | 8.57E-06 | 0.029 |
| 2 | 1.83E-08 | 1.45E-02 | -0.0144988 | -1.6214064 | 1.95E-02 | 7.62E-06 | 0.058 |
| 3 | 4.34E-06 | 9.78E-02 | -0.0977583 | -5.1346023 | 1.14E+00 | 9.80E-06 | 0.063 |
| 4 | 3.96E-04 | 3.70E-01 | -0.369611 | -6.4914696 | 3.03E+00 | 1.37E-05 | 0.124 |

| | | | | | | | |
|---|---|---|---|---|---|---|---|
| **5** | 1.63E-02 | 9.70E-01 | -0.9537708 | -10.329422 | 1.10E+01 | 1.71E-05 | 0.242 |
| **6** | 3.30E-01 | 2.13E+00 | -1.7997457 | -12.339504 | 3.01E+01 | 2.29E-05 | 0.491 |
| **7** | 3.39E+00 | 4.36E+00 | -0.9752849 | -12.008474 | 4.91E+01 | 6.86E-05 | 0.893 |
| **8** | 1.79E+01 | 7.93E+00 | 10.0101164 | -13.003822 | 6.48E+01 | 3.43E-05 | 0.588 |
| **9** | 5.05E+01 | 1.16E+01 | 38.8497452 | -13.378246 | 7.40E+01 | 1.14E-05 | 0.1285 |

##### Supplementary Table 25. Dark Glial Cell Density POL L vs POL E Females difference

| Decile | POL L Females | POL E Females | Difference | CI lower | CI upper | p crit | p-value |
|---|---|---|---|---|---|---|---|
| **1** | 2.07428 | 2.67E-07 | 2.07428 | 0.00506411 | 20.39504 | 1.14E-05 | <0.001 |
| **2** | 5.177012 | 1.77E-05 | 5.176995 | 0.24945673 | 21.60263 | 5.72E-06 | <0.001 |
| **3** | 9.1135 | 4.38E-04 | 9.113062 | 1.58581475 | 27.90209 | 3.81E-06 | <0.001 |
| **4** | 14.175393 | 5.66E-03 | 14.169732 | 3.19963492 | 29.24835 | 2.86E-06 | <0.001 |
| **5** | 19.610566 | 4.37E-02 | 19.566873 | 4.35266336 | 30.91274 | 2.29E-06 | <0.001 |
| **6** | 24.351703 | 2.17E-01 | 24.134496 | 6.61876696 | 38.23302 | 1.91E-06 | <0.001 |
| **7** | 28.086174 | 7.34E-01 | 27.35232 | 8.18192076 | 43.75649 | 1.63E-06 | <0.001 |
| **8** | 32.027146 | 1.77E+00 | 30.256825 | 12.0509654 | 49.0833 | 1.43E-06 | <0.001 |
| **9** | 40.542751 | 3.17E+00 | 37.370737 | 19.7019744 | 49.96782 | 1.27E-06 | <0.001 |

#### Supplementary Table 26. Dark Neuron Density POL E vs SAL Males difference

| Decile | POL E Males | SAL Males | Difference | CI lower | CI upper | p crit | p-value |
|---|---|---|---|---|---|---|---|
| 1 | 7.95E-07 | 0.01338757 | -0.0133868 | -1.656836 | 0.01105213 | 0.00555556 | 0.087 |
| 2 | 5.23E-05 | 0.21362932 | -0.2135771 | -6.000844 | 0.15721379 | 0.00277778 | 0.053 |
| 3 | 1.28E-03 | 1.12113573 | -1.1198526 | -6.815987 | 0.7895302 | 0.00185185 | 0.06 |
| 4 | 1.63E-02 | 2.94651928 | -2.9302176 | -11.610068 | 0.92498822 | 0.00138889 | 0.068 |
| 5 | 1.22E-01 | 4.95042709 | -4.828388 | -21.334763 | 2.10711202 | 0.00111111 | 0.037 |
| 6 | 5.76E-01 | 6.83510042 | -6.2595173 | -26.797794 | 2.27674329 | 0.00092593 | 0.024 |
| 7 | 1.78E+00 | 10.0453915 | -8.2656551 | -32.964434 | 1.62940028 | 0.00079365 | 0.014 |
| 8 | 3.71E+00 | 17.3375998 | -13.625091 | -33.323013 | 0.00326122 | 0.00069444 | 0.003 |
| 9 | 5.41E+00 | 29.53897 | -24.128274 | -36.842743 | -0.3306611 | 0.00061728 | 0.001 |

#### Supplementary Table 27. Dark Neuron Density POL E vs SAL Females difference

| Decile | POL E Females | SAL Females | Difference | CI lower | CI upper | p crit | p-value |
|---|---|---|---|---|---|---|---|
| 1 | 1.11E-05 | 1.63E-09 | 1.11E-05 | -0.0093335 | 0.1719587 | 8.82E-05 | 0.317 |
| 2 | 4.38E-04 | 9.10E-07 | 4.37E-04 | -0.0929765 | 0.5793316 | 1.03E-04 | 0.357 |
| 3 | 6.47E-03 | 1.04E-04 | 6.36E-03 | -0.6910162 | 1.7857513 | 1.23E-04 | 0.483 |
| 4 | 4.97E-02 | 3.99E-03 | 4.57E-02 | -2.3319386 | 1.9669653 | 1.54E-04 | 0.609 |
| 5 | 2.28E-01 | 6.49E-02 | 1.63E-01 | -3.6154481 | 2.3426595 | 3.09E-04 | 0.744 |
| 6 | 6.77E-01 | 5.07E-01 | 1.69E-01 | -6.5124234 | 2.710026 | 6.17E-04 | 0.963 |

| | | | | | | | |
|---|---|---|---|---|---|---|---|
| 7 | 1.40E+00 | 2.10E+00 | -7.04E-01 | -8.1366172 | 2.6683049 | 2.06E-04 | 0.668 |
| 8 | 2.20E+00 | 5.11E+00 | -2.91E+00 | -10.199566 | 2.603619 | 7.72E-05 | 0.242 |
| 9 | 2.79E+00 | 8.71E+00 | -5.92E+00 | -11.88331 | 1.2480937 | 6.86E-05 | 0.019 |

#### Supplementary Table 28. Dark Neuron Density POL L vs SAL Males difference

| Decile | POL L Males | SAL Males | Difference | CI lower | CI upper | p crit | p-value |
|---|---|---|---|---|---|---|---|
| 1 | 1.80E-12 | 0.01338757 | -0.0133876 | -3.619961 | -1.57E-07 | 0.01 | 0.001 |
| 2 | 1.69E-09 | 0.21362932 | -0.2136293 | -5.673354 | -6.32E-06 | 0.00555556 | 0.002 |
| 3 | 4.07E-07 | 1.12113573 | -1.1211353 | -6.429677 | -4.73E-05 | 0.00714286 | 0.006 |
| 4 | 3.84E-05 | 2.94651928 | -2.9464809 | -9.063155 | -1.52E-03 | 0.00833333 | 0.006 |
| 5 | 1.71E-03 | 4.95042709 | -4.9487126 | -14.725415 | -1.21E-02 | 0.00625 | 0.007 |
| 6 | 4.04E-02 | 6.83510042 | -6.7947181 | -21.616597 | -3.87E-01 | 0.0125 | 0.008 |
| 7 | 5.32E-01 | 10.0453915 | -9.5133155 | -27.359307 | 3.69E+00 | 0.01666667 | 0.053 |
| 8 | 3.85E+00 | 17.3375998 | -13.483337 | -31.442905 | 5.99E+00 | 0.025 | 0.132 |
| 9 | 1.40E+01 | 29.53897 | -15.579848 | -31.565599 | 6.54E+00 | 0.05 | 0.177 |

#### Supplementary Table 29. Dark Neuron Density POL L vs SAL Females difference

| Decile | POL L Females | SAL Females | Difference | CI lower | CI upper | p crit | p-value |
|---|---|---|---|---|---|---|---|
| 1 | 1.32E-04 | 1.63E-09 | 0.00013228 | -0.0002651 | 0.3096555 | 0.00625 | 0.136 |
| 2 | 5.25E-03 | 9.10E-07 | 0.00524855 | -0.0151697 | 1.7952565 | 0.01 | 0.159 |
| 3 | 7.19E-02 | 1.04E-04 | 0.07180021 | -0.1671349 | 5.7450419 | 0.00833333 | 0.198 |

| | | | | | | | |
|---|---|---|---|---|---|---|---|
| **4** | 4.91E-01 | 3.99E-03 | 0.4866887 | -0.6373064 | 8.9532267 | 0.0125 | 0.206 |
| **5** | 1.94E+00 | 6.49E-02 | 1.87750889 | -0.6823101 | 8.6566387 | 0.05 | 0.206 |
| **6** | 4.86E+00 | 5.07E-01 | 4.35560012 | -1.9815046 | 10.6758468 | 0.025 | 0.190 |
| **7** | 8.32E+00 | 2.10E+00 | 6.21590361 | -4.3655219 | 12.1094618 | 0.01666667 | 0.202 |
| **8** | 1.11E+01 | 5.11E+00 | 5.97530913 | -6.2419107 | 14.2211203 | 0.00714286 | 0.129 |
| **9** | 1.43E+01 | 8.71E+00 | 5.60573642 | -5.7310481 | 14.0907629 | 0.00555556 | 0.118 |

#### Supplementary Table 30. Dark Neuron Density POL L vs POL E Males difference

| Decile | POL L Males | POL E Males | Difference | CI lower | CI upper | p crit | p-value |
|---|---|---|---|---|---|---|---|
| **1** | 1.80E-12 | 7.95E-07 | -7.95E-07 | -0.0928224 | 3.90E-05 | 9.80E-06 | 0.236 |
| **2** | 1.69E-09 | 5.23E-05 | -5.23E-05 | -0.7129505 | 1.96E-03 | 8.57E-06 | 0.3375 |
| **3** | 4.07E-07 | 1.28E-03 | -1.28E-03 | -2.2161713 | 1.00E-01 | 1.14E-05 | 0.314 |
| **4** | 3.84E-05 | 1.63E-02 | -1.63E-02 | -3.156025 | 7.54E-01 | 1.37E-05 | 0.357 |
| **5** | 1.71E-03 | 1.22E-01 | -1.20E-01 | -5.8132401 | 3.74E+00 | 1.71E-05 | 0.442 |
| **6** | 4.04E-02 | 5.76E-01 | -5.35E-01 | -5.6806593 | 1.05E+01 | 2.29E-05 | 0.551 |
| **7** | 5.32E-01 | 1.78E+00 | -1.25E+00 | -5.7970739 | 1.56E+01 | 3.43E-05 | 0.700 |
| **8** | 3.85E+00 | 3.71E+00 | 1.42E-01 | -5.9888879 | 1.89E+01 | 6.86E-05 | 0.937 |
| **9** | 1.40E+01 | 5.41E+00 | 8.55E+00 | -5.996452 | 2.20E+01 | 7.62E-06 | 0.2705 |

#### Supplementary Table 31. Dark Neuron Density POL L vs POL E Females difference

| Decile | POL L Females | POL E Females | Difference | CI lower | CI upper | p crit | p-value |
|---|---|---|---|---|---|---|---|
| 1 | 1.32E-04 | 1.11E-05 | 1.21E-04 | -0.3745774 | 1.694192 | 7.62E-06 | 0.663 |
| 2 | 5.25E-03 | 4.38E-04 | 4.81E-03 | -0.5793341 | 2.711498 | 3.81E-06 | 0.556 |
| 3 | 7.19E-02 | 6.47E-03 | 6.54E-02 | -1.8740314 | 8.01383 | 2.54E-06 | 0.510 |
| 4 | 4.91E-01 | 4.97E-02 | 4.41E-01 | -1.836736 | 10.340975 | 1.91E-06 | 0.459 |
| 5 | 1.94E+00 | 2.28E-01 | 1.71E+00 | -2.3481622 | 11.328048 | 1.52E-06 | 0.331 |
| 6 | 4.86E+00 | 6.77E-01 | 4.19E+00 | -2.3535711 | 12.426122 | 1.27E-06 | 0.183 |
| 7 | 8.32E+00 | 1.40E+00 | 6.92E+00 | -2.7252112 | 13.838796 | 1.09E-06 | 0.084 |
| 8 | 1.11E+01 | 2.20E+00 | 8.88E+00 | -2.5829455 | 16.457213 | 9.53E-07 | 0.015 |
| 9 | 1.43E+01 | 2.79E+00 | 1.15E+01 | -2.4035657 | 17.33054 | 8.47E-07 | 0.006 |

#### Supplementary Table 32. Apoptotic Cell Density POL E vs SAL Males difference

| Decile | POL E Males | SAL Males | Difference | CI lower | CI upper | p crit | p-value |
|---|---|---|---|---|---|---|---|
| 1 | 5.75E-06 | 5.23E-08 | 5.70E-06 | -0.200738 | 0.08309331 | 1.32E-04 | 0.536 |
| 2 | 2.32E-04 | 7.00E-06 | 2.25E-04 | -0.1985263 | 0.52506395 | 1.13E-04 | 0.529 |
| 3 | 3.57E-03 | 2.89E-04 | 3.29E-03 | -0.7255787 | 1.17588146 | 1.59E-04 | 0.534 |
| 4 | 2.94E-02 | 5.34E-03 | 2.41E-02 | -1.3666148 | 2.05924476 | 2.65E-04 | 0.644 |
| 5 | 1.51E-01 | 5.09E-02 | 1.00E-01 | -1.8282966 | 3.81782995 | 7.94E-04 | 0.715 |

| | | | | | | | |
|---|---|---|---|---|---|---|---|
| 6 | 5.41E-01 | 2.70E-01 | 2.71E-01 | -2.5794743 | 5.62691195 | 3.97E-04 | 0.670 |
| 7 | 1.47E+00 | 8.33E-01 | 6.41E-01 | -3.0073851 | 6.51577473 | 1.98E-04 | 0.645 |
| 8 | 3.27E+00 | 1.65E+00 | 1.63E+00 | -3.0865078 | 6.58912238 | 9.92E-05 | 0.443 |
| 9 | 5.94E+00 | 2.73E+00 | 3.21E+00 | -3.799139 | 7.4130836 | 8.82E-05 | 0.279 |

#### Supplementary Table 33. Apoptotic Cell Density POL E vs SAL Females difference

| Decile | POL E Females | SAL Females | Difference | CI lower | CI upper | p crit | p-value |
|---|---|---|---|---|---|---|---|
| 1 | 0.06077409 | 2.19E-08 | 0.06077406 | -5.89E-05 | 2.003436 | 9.80E-06 | 0.001 |
| 2 | 0.32815958 | 7.40E-06 | 0.32815218 | -4.91E-02 | 2.109396 | 1.10E-05 | 0.005 |
| 3 | 0.80343016 | 5.01E-04 | 0.80292952 | -5.25E-01 | 2.564185 | 1.26E-05 | 0.011 |
| 4 | 1.33231219 | 1.12E-02 | 1.3210813 | -6.16E-01 | 2.998568 | 1.47E-05 | 0.011 |
| 5 | 1.81378203 | 1.05E-01 | 1.70897342 | -6.95E-01 | 4.165108 | 1.76E-05 | 0.020 |
| 6 | 2.28556947 | 4.73E-01 | 1.81283812 | -1.20E+00 | 4.706171 | 2.20E-05 | 0.029 |
| 7 | 2.8913935 | 1.17E+00 | 1.72067703 | -1.03E+00 | 4.483863 | 2.94E-05 | 0.044 |
| 8 | 3.74678314 | 1.86E+00 | 1.884857 | -1.68E+00 | 4.577465 | 4.41E-05 | 0.064 |
| 9 | 4.6245176 | 2.88E+00 | 1.74712351 | -1.97E+00 | 4.056836 | 8.82E-05 | 0.168 |

#### Supplementary Table 34. Apoptotic Density POL L vs SAL Males difference

| Decile | POL L Males | SAL Males | Difference | CI lower | CI upper | p crit | p-value |
|---|---|---|---|---|---|---|---|
| 1 | 3.28E-09 | 5.23E-08 | -4.90E-08 | -0.0044623 | 0.00031029 | 0.00625 | 0.787 |

| | | | | | | | |
|---|---|---|---|---|---|---|---|
| **2** | 9.07E-07 | 7.00E-06 | -6.10E-06 | -0.071187 | 0.0107912 | 0.00833333 | 0.794 |
| **3** | 6.39E-05 | 2.89E-04 | -2.25E-04 | -0.0538405 | 0.03489436 | 0.05 | 0.844 |
| **4** | 1.75E-03 | 5.34E-03 | -3.58E-03 | -0.5735756 | 0.21625223 | 0.01666667 | 0.823 |
| **5** | 2.22E-02 | 5.09E-02 | -2.87E-02 | -1.5118993 | 0.77329311 | 0.0125 | 0.830 |
| **6** | 1.45E-01 | 2.70E-01 | -1.25E-01 | -1.6858897 | 1.0409658 | 0.025 | 0.878 |
| **7** | 5.32E-01 | 8.33E-01 | -3.01E-01 | -2.3399603 | 1.59191438 | 0.01 | 0.763 |
| **8** | 1.25E+00 | 1.65E+00 | -3.94E-01 | -2.8467626 | 2.50316015 | 0.00555556 | 0.718 |
| **9** | 2.41E+00 | 2.73E+00 | -3.13E-01 | -2.8257589 | 3.08950572 | 0.00714286 | 0.766 |

**Supplementary Table 35. Apoptotic Cell Density POL L vs SAL Females difference**

| Decile | POL L Females | SAL Females | Difference | CI lower | CI upper | p crit | p-value |
|---|---|---|---|---|---|---|---|
| **1** | 3.56E-09 | 2.19E-08 | -1.84E-08 | -0.0012782 | 0.0005374 | 0.00714286 | 0.809 |
| **2** | 5.54E-07 | 7.40E-06 | -6.84E-06 | -0.0885355 | 0.02471114 | 0.00357143 | 0.683 |
| **3** | 2.84E-05 | 5.01E-04 | -4.72E-04 | -0.461889 | 0.15471194 | 0.00238095 | 0.601 |
| **4** | 6.82E-04 | 1.12E-02 | -1.05E-02 | -0.9482446 | 0.42394535 | 0.00178571 | 0.551 |
| **5** | 8.92E-03 | 1.05E-01 | -9.59E-02 | -1.7132934 | 0.96299854 | 0.00142857 | 0.494 |
| **6** | 6.83E-02 | 4.73E-01 | -4.04E-01 | -2.2593282 | 1.17609701 | 0.00119048 | 0.399 |
| **7** | 3.19E-01 | 1.17E+00 | -8.52E-01 | -2.4625073 | 1.39261982 | 0.00102041 | 0.334 |
| **8** | 9.27E-01 | 1.86E+00 | -9.35E-01 | -3.5729426 | 1.37456302 | 0.00089286 | 0.252 |
| **9** | 1.68E+00 | 2.88E+00 | -1.20E+00 | -4.5062637 | 0.9738635 | 0.00079365 | 0.084 |

**Supplementary Table 36. Apoptotic Cell Density POL L vs POL E Males difference**

| Decile | POL L Males | POL E Males | Difference | CI lower | CI upper | p crit | p-value |
|---|---|---|---|---|---|---|---|
| 1 | 3.28E-09 | 5.75E-06 | -5.75E-06 | -0.1954404 | 0.00559849 | 1.10E-05 | 0.338 |
| 2 | 9.07E-07 | 2.32E-04 | -2.31E-04 | -2.004306 | 0.03225349 | 1.47E-05 | 0.373 |
| 3 | 6.39E-05 | 3.57E-03 | -3.51E-03 | -2.3097949 | 0.37140159 | 1.76E-05 | 0.412 |
| 4 | 1.75E-03 | 2.94E-02 | -2.77E-02 | -3.1251772 | 0.86786421 | 2.94E-05 | 0.505 |
| 5 | 2.22E-02 | 1.51E-01 | -1.29E-01 | -3.8677683 | 0.96845351 | 8.82E-05 | 0.528 |
| 6 | 1.45E-01 | 5.41E-01 | -3.96E-01 | -6.1913104 | 1.35167046 | 4.41E-05 | 0.548 |
| 7 | 5.32E-01 | 1.47E+00 | -9.41E-01 | -7.5731441 | 1.73926306 | 2.20E-05 | 0.449 |
| 8 | 1.25E+00 | 3.27E+00 | -2.02E+00 | -7.7372568 | 2.34876766 | 1.26E-05 | 0.344 |
| 9 | 2.41E+00 | 5.94E+00 | -3.53E+00 | -7.6425497 | 3.35488955 | 9.80E-06 | 0.248 |

**Supplementary Table 37. Apoptotic Cell Density POL L vs POL E Females difference**

| Decile | POL L Females | POL E Females | Difference | CI lower | CI upper | p crit | p-value |
|---|---|---|---|---|---|---|---|
| 1 | 3.56E-09 | 0.06077409 | -0.0607741 | -2.016653 | 2.90E-03 | 1.09E-06 | 0.002 |
| 2 | 5.54E-07 | 0.32815958 | -0.328159 | -2.439921 | -2.59E-05 | 1.96E-06 | <0.001 |
| 3 | 2.84E-05 | 0.80343016 | -0.8034018 | -2.835063 | 9.02E-03 | 4.90E-06 | 0.002 |
| 4 | 6.82E-04 | 1.33231219 | -1.3316299 | -3.844567 | -6.84E-03 | 3.27E-06 | 0.001 |
| 5 | 8.92E-03 | 1.81378203 | -1.8048616 | -4.413767 | 3.18E-01 | 1.63E-06 | 0.005 |

| | | | | | | | |
|---|---|---|---|---|---|---|---|
| **6** | 6.83E-02 | 2.28556947 | -2.217292 | -4.535999 | 1.47E-01 | 2.45E-06 | 0.003 |
| **7** | 3.19E-01 | 2.8913935 | -2.5723796 | -4.869634 | 8.07E-01 | 1.40E-06 | 0.005 |
| **8** | 9.27E-01 | 3.74678314 | -2.8199705 | -4.978101 | -1.08E-01 | 9.80E-06 | 0.001 |
| **9** | 1.68E+00 | 4.6245176 | -2.9462621 | -4.993 | -8.56E-02 | 1.22E-06 | <0.001 |
